# Multi-omics and biochemical reconstitution reveal CDK7-dependent mechanisms controlling RNA polymerase II function at gene 5’- and 3’-ends

**DOI:** 10.1101/2025.01.08.632016

**Authors:** Olivia Luyties, Lynn Sanford, Jessica Rodino, Michael Nagel, Taylor Jones, Jenna K. Rimel, Christopher C. Ebmeier, Grace S. Shelby, Kira Cozzolino, Finn Brennan, Axel Hartzog, Mirzam B. Saucedo, Lotte P. Watts, Sabrina Spencer, Jennifer F. Kugel, Robin D. Dowell, Dylan J. Taatjes

**Affiliations:** Dept. of Biochemistry, University of Colorado; Boulder, CO, 80303, USA; Dept. of Molecular, Cellular, and Developmental Biology, University of Colorado; Boulder, CO, 80303, USA; BioFrontiers Institute, University of Colorado; Boulder, CO, 80303, USA

## Abstract

CDK7 regulates RNA polymerase II (RNAPII) initiation, elongation, and termination through incompletely understood mechanisms. Because contaminating kinases precluded CDK7 analysis with nuclear extracts, we completed biochemical assays with purified factors. Reconstitution of RNAPII transcription initiation showed CDK7 inhibition slowed and/or paused RNAPII promoter-proximal transcription, which reduced re-initiation. These CDK7-regulatory functions were Mediator- and TFIID-dependent. Similarly in human cells, CDK7 inhibition reduced transcription by suppressing RNAPII activity at promoters, consistent with reduced initiation and/or re-initiation. Moreover, widespread 3’-end readthrough transcription was observed in CDK7-inhibited cells; mechanistically, this occurred through rapid nuclear depletion of RNAPII elongation and termination factors, including high-confidence CDK7 targets. Collectively, these results define how CDK7 governs RNAPII function at gene 5’-ends and 3’-ends, and reveal that nuclear abundance of elongation and termination factors is kinase-dependent. Because 3’-readthrough transcription is commonly induced during stress, our results further suggest regulated suppression of CDK7 activity may enable this RNAPII transcriptional response.

## INTRODUCTION

The human Pre-Initiation Complex (PIC) consists of RNA polymerase II (RNAPII), TFIIA, TFIIB, TFIID, TFIIE, TFIIF, TFIIH, and Mediator.^1^ The PIC structure reveals that the 10-subunit TFIIH complex assembles and orients through interactions with Mediator, TFIID, and promoter DNA.^2-4^ TFIIH performs multiple key functions during transcription initiation. For example, the XPB helicase helps open the DNA template at the transcription start site (TSS), allowing single-stranded DNA to enter the RNAPII active site. CDK7 is part of the TFIIH kinase module, which interacts with Mediator in the PIC.^2,3^ In its unphosphorylated form, the RNAPII C-terminal domain (CTD) interacts with Mediator,^5,6^ and this interaction guides the structurally disordered CTD to the CDK7 active site.^2,3^ CTD phosphorylation by CDK7, which occurs during RNAPII transcription initiation,^7^ disrupts the RNAPII CTD-Mediator interaction based upon biochemical and proteomics data.^8-10^ CDK7-dependent disruption of RNAPII CTD-Mediator interactions may enhance RNAPII initiation and promoter escape, but this hasn’t been directly tested.

The RNAPII CTD has a consensus sequence Y_1_S_2_P_3_T_4_S_5_P_6_S_7_ repeated 26 times in *S. cerevisiae* and 52 times in humans.^11^ CDK7 (or its *S. cerevisiae* ortholog Kin28) phosphorylates the RNAPII CTD at ser5 and ser7 positions within the CTD heptad repeat. Specific protein factors recognize distinct CTD phosphorylation marks, and this helps regulate the timing of co-transcriptional events such as capping, splicing, 3’-end cleavage, termination, and poly-adenylation.^11^

A number of studies have used chemical inhibitors or chemical genetics to assess how CDK7 function affects RNAPII transcription in cells. In general, these experiments measured steady-state mRNA levels after 4-24h inhibitor treatment. In yeast, Kin28 inhibition had little effect on RNAPII occupancy at gene 5’-ends (as measured by ChIP-PCR or ChIP-seq); however, RNAPII CTD phosphorylation was disrupted at ser5 and ser7.^12^ Upon Kin28 inhibition in yeast, nascent RNAPII transcription and mRNA biogenesis were reduced; however, this was offset by mRNA stabilization, such that overall mRNA levels remained largely unchanged within the timeframes of the experiments.^13^

In human cells, RNA-seq data showed both up- and down-regulation of hundreds of genes 5-24h after CDK7 inhibition.^8,14,15^ Furthermore, CDK7 inhibition broadly and rapidly (within 30min) suppresses RNAPII transcriptional output, as shown by TT-seq analysis of nascent transcription.^16^ CDK7 inhibition also changes the distribution of RNAPII at gene 5’-ends; in particular, RNAPII promoter-proximal pausing increased based upon ChIP-seq experiments.^8^ More recent work showed evidence for RNAPII inactivation at promoters in CDK7-inhibited cells, with RNAPII accumulation upstream of the transcription start site.^16^ These intriguing results demonstrate that CDK7 kinase activity affects RNAPII initiation at gene 5’-ends, but the underlying mechanisms remain incompletely understood. ChIP and ChIP-seq data also showed evidence for readthrough defects at gene 3’-ends in CDK7-inhibited cells,^8,17,18^ but here as well, a mechanistic basis is lacking.

In this study, we combine biochemical reconstitution, chemical genetics, quantitative proteomics and phosphoproteomics, transcriptomics, and other methods to better understand how CDK7 kinase activity controls RNAPII function at gene 5’-ends and gene 3’-ends. We demonstrate that CDK7 inhibition slows and/or pauses RNAPII transcription through the promoter-proximal region, which collectively acts to suppress re-initiation. Separately, we discover that RNAPII elongation and termination factors are rapidly depleted upon CDK7 inhibition, revealing a simple mechanism for readthrough defects at gene 3’-ends. These distinct effects collectively provide a mechanistic basis for reduced RNAPII activity at gene 5’-ends and dysregulated transcription at gene 3’-ends in CDK7-inhibited cells.

## RESULTS

### Nuclear extracts are poorly suited to assess human CDK7 function in transcription

Analysis of RNAPII transcription *in vitro* allows more precise experimental control compared with a population of proliferating cells, for a better assessment of molecular mechanisms. A limitation with reconstituted RNAPII transcription with purified components is that it doesn’t match the complexity of factors that converge on active genes in a living cell. Nuclear extracts can be used to bridge this gap, but nuclear extracts have their own drawbacks, including a complex and undefined composition of proteins and nucleic acids. Prior work by us^7^ and others^19,20^ have shown that PICs will assemble if promoter DNA is incubated with nuclear extract.

To test whether nuclear extracts might be helpful to study CDK7-dependent mechanisms of RNAPII initiation, we immobilized the native HSPA1B promoter and incubated it with nuclear extracts prepared from HeLa cells (**Figure 1A**). After washing to remove unbound factors, we detected all PIC factors (TFIIA, TFIIB, TFIID, TFIIE, TFIIF, TFIIH, Mediator, and RNAPII) bound to the promoter, as expected (**Figure 1B**). We then added NTPs to initiate transcription, and this was done in the presence/absence of the potent and selective CDK7 inhibitor SY-5609 ^21^. As a control, we also ran experiments with flavopiridol, a pan-kinase inhibitor that targets CDK7, CDK8, CDK9, CDK12, and CDK13.^22-24^ To test whether RNAPII CTD phosphorylation was blocked in SY-5609-treated samples, we probed RNAPII by western blot. Contrary to expectations, RNAPII phosphorylation levels remained largely unchanged ±SY-5609 or ±SY-351 (**Figure 1C, D**), which is a structurally distinct covalent CDK7 inhibitor;^15^ furthermore, we observed RNAPII CTD phosphorylation even in the presence of flavopiridol, suggesting “contaminating” activity from other kinases in the extract. (We separately confirmed that SY-5609 effectively blocks CDK7 activity *in vitro*, see below.)

**Figure 1.**
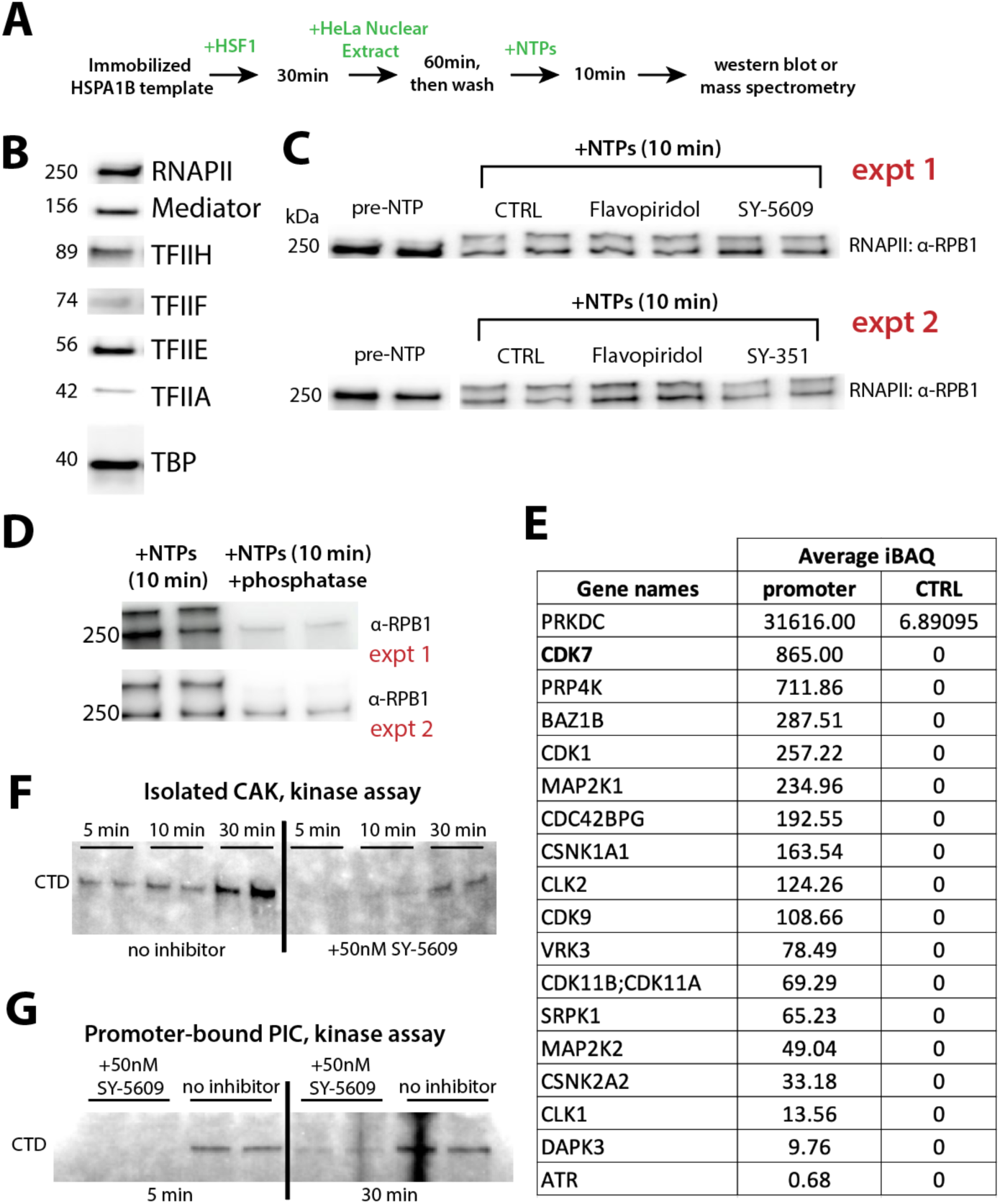
Biochemical reconstitution can specifically address CDK7-dependent functions, whereas nuclear extracts cannot. (A) Overview of immobilized template experiments. (B) Western blots showing all PIC factors bound to the HSPA1B promoter template, as expected. (C) RNAPII western blots show shifted band following NTP addition, consistent with RPB1 CTD phosphorylation. Technical replicates shown side-by-side for each condition, and data from 2 different biological replicates are shown (expt 1 & expt 2). Expt 2 used a covalent CDK7 inhibitor, SY-351. (D) The shifted RPB1 band was phosphorylated, as expected, based upon incubation with λ-phosphatase. (E) Summary of kinases bound to the native HSPA1B promoter after washing, based upon mass spectrometry analysis. Although CDK7 was a top hit, many other contaminating kinases were present. The CTRL experiment used beads only (no immobilized template). (F) Purified CAK module (CDK7, CCNC, MNAT1) phosphorylates full-length mammalian GST-CTD *in vitro*; phosphorylation is blocked by 50nM SY-5609. Some phosphorylation occurred at 30min, a timeframe longer than *in vitro* transcription reactions. Note ^56^ is 20,000-fold higher than [SY-5609] in these experiments (1mM vs. 50nM). (G) Purified reconstituted PICs assembled on the HSPA1B promoter contain only one kinase, CDK7. These PICs phosphorylate the RNAPII CTD, as expected, but phosphorylation is blocked by SY-5609, although some phosphorylation can be detected after 30min.

The human proteome contains over 500 kinases,^25^ and the RNAPII CTD contains 237 phosphorylatable residues. To identify kinases from the nuclear extract that remained bound to the HSPA1B promoter after washing, we next completed mass spectrometry experiments. The data (**Table S1**) showed the presence of CDK7 along with 17 “contaminating” kinases, including many known to modify the RNAPII CTD (**Figure 1E**). Based upon these experiments, we conclude that extracts cannot reliably assess CDK7-dependent regulation of RNAPII function, although we emphasize that *in vitro* experiments with extracts can be valuable for other purposes.

### CDK7 enhances promoter-proximal RNAPII transcription to facilitate re-initiation

We previously established an *in vitro* transcription system reconstituted from purified human PIC factors.^26^ The PIC (**Figure 2A**) contains only one kinase, CDK7, which allowed us to specifically assess its contributions to RNAPII initiation and promoter-proximal pausing using the well-studied human HSPA1B promoter. An overview of the “pausing assay” is shown in **Figure 2B**. A key feature of this assay is a pulse-chase protocol in which transcription is initiated by adding ATP, GTP, UTP, and ^32^P-CTP for 1 minute followed by a 4-minute “chase” with addition of a high concentration of non-radioactive CTP (5 min reaction); alternatively, reactions were immediately stopped (1 min reaction). Because of the initial ^32^P-CTP pulse, short transcripts can be visualized despite fewer incorporated ^32^P-cytidines relative to longer transcripts. The initial pulse also favors labeling of first-round RNAPII transcripts compared with transcripts that result from re-initiation events (e.g. from a second RNAPII enzyme). The timeframes for these assays were determined empirically; we observed that CDK7-dependent effects occurred within 5 minutes, presumably due to secondary effects from CDK7 inhibition at longer timeframes (**Figure S1A**). Furthermore, *in vitro* kinase assays showed that 50nM SY-5609 completely blocked RNAPII CTD phosphorylation within the 5min timeframe (**Figure 1F, G**). We therefore focused our experiments within 5min NTP addition. An additional rationale for short timeframes was that in cells, RNAPII transcription occurs during short bursts of multi-round initiation events, followed by prolonged dormant periods.^27^

**Figure 2.**
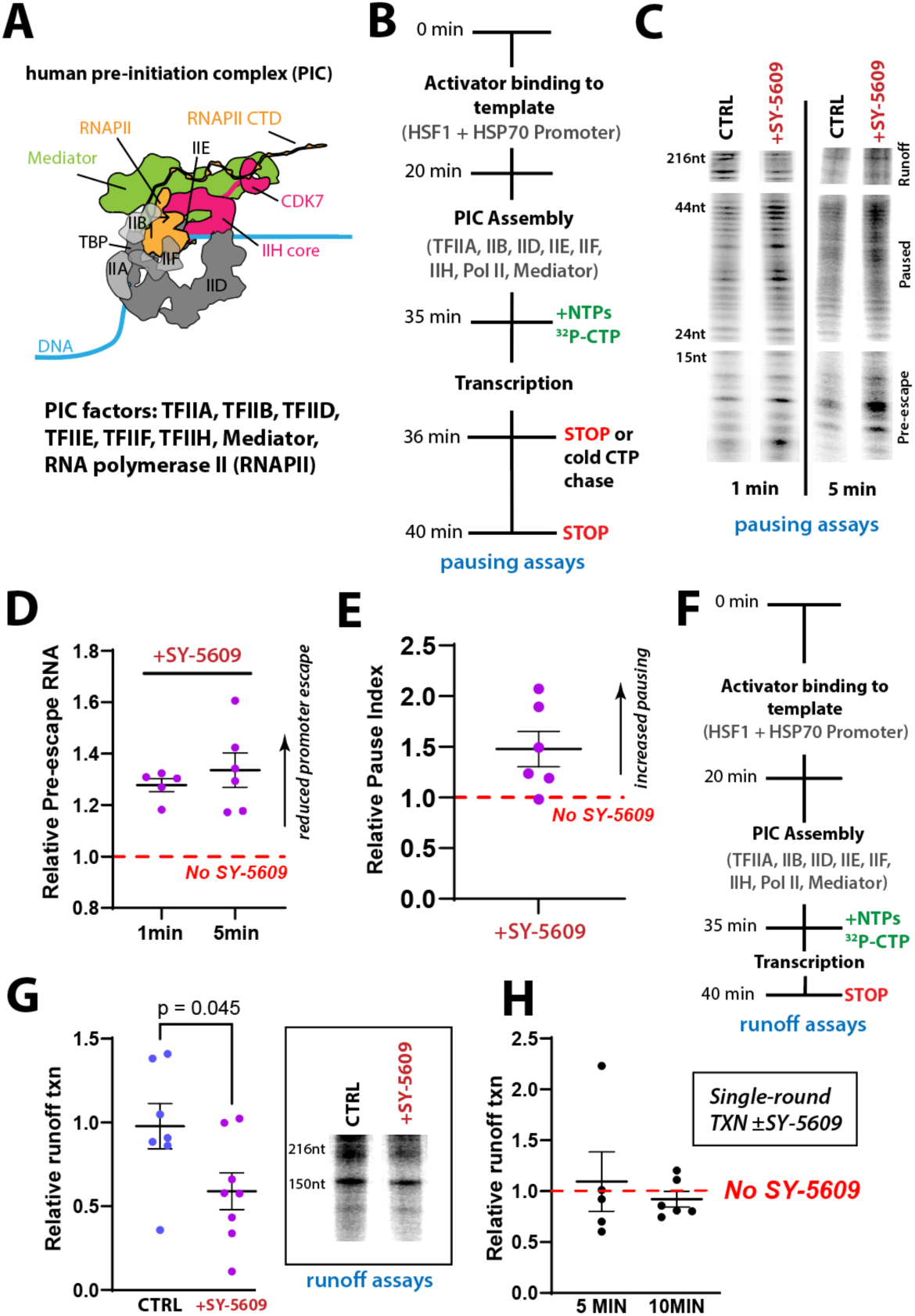
CDK7 inhibition blocks promoter escape and pause release, and reduces overall transcription output. (A) Schematic of the human pre-initiation complex (PIC). (B) Overview of the *in vitro* transcription pausing assays. (C) Representative gel images of pausing assays, with pre-escape, paused, and runoff/elongated regions indicated. Each region was independently contrast-enhanced to aid visualization and each region was adjusted equally across comparisons (e.g. CTRL vs. SY-5609). Quantitation was completed with uniform exposure across the entire lane; gel lanes with uniform exposure are shown in **Figure S1B**. (D) CDK7 inhibition decreases and slows promoter escape. Each point (n=6) represents the ratio of pre-escape transcripts (5-15nt length) to runoff products (150-216nt) from reactions treated with 50nM SY-5609, normalized to no inhibitor controls. (E) CDK7 inhibition increases pausing. Plot shows the pause index of reactions (n=6) treated with SY-5609, normalized to no inhibitor controls. (F) Overview of *in vitro* transcription runoff assays. (G) CDK7 inhibition reduces overall RNAPII transcriptional output (n=7). Inset: representative gel images of runoff transcription ±SY-5609. See also **Figure S1C**. (H) CDK7 inhibition under single-round conditions (sarkosyl added 1-min post-NTP; n=5). Plot shows quantification of runoff transcription +SY-5609, normalized to no inhibitor controls. Representative gels shown in **Figure S1D**.

As shown in **Figure 2C**, the pausing assays allowed detection of 3 distinct sets of transcripts: promoter-associated (prior to promoter escape), promoter-proximal paused, or elongated “runoff” transcripts. (Note that the assay cannot distinguish released transcripts from those that remain RNAPII-associated; full uniform-contrast lanes in **Figure S1B**.) As shown in **Figure 2C-E**, CDK7 inhibition (50nM SY-5609) increased transcript levels in the promoter-associated and paused regions during the 5-minute pulse-chase experiments. We also probed RNAPII transcription after only 1min NTP addition to obtain a “snapshot” of actively transcribing complexes. The data were consistent with the 5-minute pulse-chase results (**Figure 2C, D**). Although a caveat is that many RNAPII complexes will be actively transcribing through the pause region at the 1-minute timepoint, these data suggest that CDK7 kinase activity accelerates RNAPII promoter escape and pause release in this reconstituted PIC system. We emphasize that this system lacks post-initiation regulatory factors such as NELF and DSIF; thus, the PIC alone is sufficient for these CDK7-dependent effects.

To further test how CDK7 activity impacted overall transcriptional output, we completed a series of “runoff” assays (**Figure 2F**). Unlike the pausing assays, these experiments did not implement a pulse-chase but instead involved a single NTP addition step (ATP, GTP, UTP, and ^32^P-CTP + non-radioactive CTP). Although short transcripts were harder to visualize in these assays, the consistent concentration of ^32^P-CTP throughout the experiment allowed a more reliable quantitation of runoff transcripts, including those that resulted from RNAPII re-initiation events. As shown in **Figure 2G**, the runoff assays showed that CDK7 inhibition reduced RNAPII transcriptional output by about 40% compared with controls (see also **Figure S1C**).

Because a promoter-associated RNAPII complex, including paused RNAPII, will physically block re-initiation by another RNAPII enzyme,^28,29^ the results summarized in **Figure 2A-G** were consistent with reduced re-initiation upon CDK7 inhibition. However, it remained possible that fewer PICs successfully initiated RNAPII transcription in the absence of CDK7 activity. To test this idea, we compared CDK7-dependent RNAPII transcriptional output under single-vs. multi-round conditions. The Roeder lab has shown that the addition of sarkosyl will prevent RNAPII re-initiation while allowing promoter-bound PICs and/or transcriptionally engaged RNAPII to transcribe normally.^30^ As shown in **Figure 2H** (see also **Figure S1D**), single-round transcription was not negatively affected by CDK7 inhibition, in contrast with RNAPII transcriptional output under standard “multi-round” runoff assay conditions (i.e. no sarkosyl) that allow re-initiation to occur. A separate set of single-round sarkosyl-treated experiments was completed with a 10-minute timeframe to ensure that all first-round runoff transcripts were detected (**Figure 2H**), and control experiments confirmed that more transcripts were generated from multi-round (no sarkosyl) vs. single-round experiments (+sarkosyl), as expected (**Figure S1E**).

Collectively, the data summarized in **Figure 2** reveal that CDK7 inhibition reduced overall RNAPII transcriptional output. Mechanistically, this occurred through slowing and/or pausing promoter-proximal RNAPII transcription. However, reduced output was observed only under multi-round conditions; single-round output was similar ±CDK7 inhibition. Because multi-round transcription requires re-initiation, we conclude that CDK7 kinase activity promotes RNAPII re-initiation. This model is also generally consistent with a study using CDK7 analog-sensitive (CDK7AS) cells.^16^ To further assess potential mechanisms, we tested whether Mediator and/or TFIID were contributing to the CDK7-dependent effects. These experiments were motivated in part by evidence that Mediator contributes to RNAPII re-initiation in cells^31^ and that TFIID regulates promoter-proximal pausing.^26^

### Mediator and TFIID coordinate CDK7-dependent regulation of promoter-proximal transcription

Among PIC factors, RNAPII interacts most extensively with Mediator,^2-4^ including through an RNAPII CTD interaction that is disrupted by CDK7-dependent CTD phosphorylation.^8-10^ The increased levels of promoter-associated transcripts upon CDK7 inhibition (e.g. **Figure 2C, D**) suggested a role for Mediator via prolonged retention of Mediator-RNAPII CTD interactions to block or delay RNAPII promoter escape. To test this hypothesis, we performed a series of multi-round “runoff” experiments in which Mediator was removed, and RNAPII transcriptional output was compared to “complete PIC” assays. As shown in **Figure 3A** (see also **Figure S1F**), the reduced transcriptional output with CDK7 inhibition was Mediator-dependent, consistent with the notion that i) Mediator regulates RNAPII promoter escape in coordination with CDK7, and that ii) retention of Mediator-RNAPII CTD interactions can block or delay RNAPII re-initiation.

**Figure 3.**
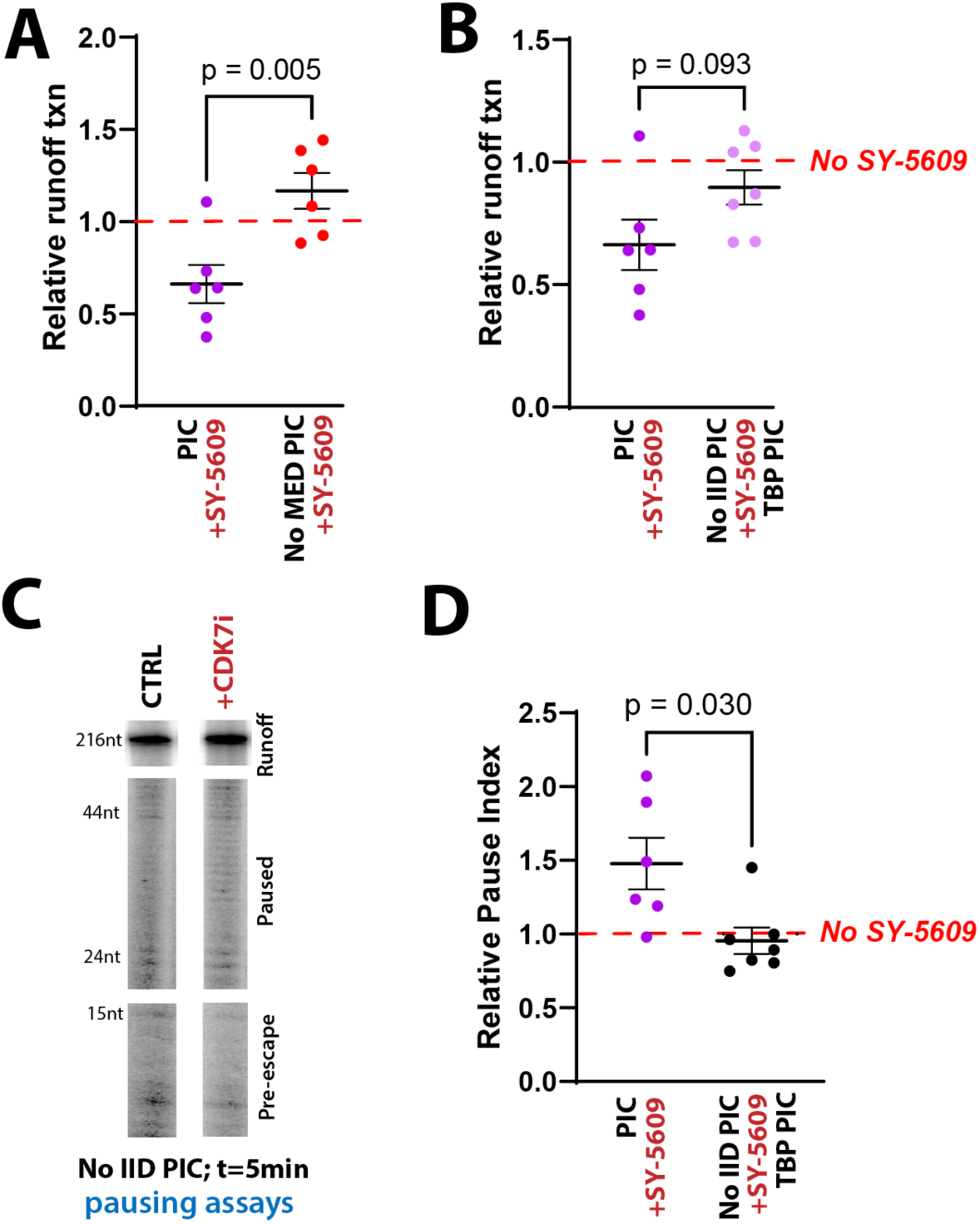
Mediator and TFIID cooperate with CDK7 to enhance promoter escape and pause release, respectively. (A) Quantitation of runoff transcription +SY-5609, normalized to no inhibitor controls (dashed line), comparing PICs with and without Mediator (n=6). Representative gel lanes in **Figure S1F**. (B) TFIID affects runoff transcription as a function of CDK7 kinase activity. Data from PICs ±TFIID shown, +SY-5609, normalized to no inhibitor controls (dashed line). No IID = TBP instead of TFIID (n=7). PIC data (left column, dark purple dots) are the same as in panel A. Representative gel lanes in **Figure S1G**. (C) Representative gel images from TBP PIC pausing assays, with pre-escape, paused, and runoff/elongated regions indicated. Each region was independently contrast-enhanced to aid visualization and each region was adjusted equally across comparisons (e.g. CTRL vs. SY-5609). Quantitation completed with uniform exposure across the entire lane; gel lanes with uniform exposure are shown in **Figure S1H**. (D) In contrast to TFIID PICs, pausing does not occur with TBP PICs and CDK7-dependent effect on pausing is lost (n=7). Purple points are the same as those in 2E, shown again here for comparison.

We previously established that the PIC (**Figure 2A**) was sufficient to establish promoter-proximal pausing and that pausing was lost if TFIID was replaced with TBP.^26^ Because RNAPII pausing increased upon CDK7 inhibition (e.g. **Figure 2E**), we hypothesized that this effect would be TFIID-dependent. Therefore, we completed multi-round runoff experiments in which TFIID was removed and replaced with TBP. Under CDK7-inhibition conditions, transcription levels increased with TBP-containing PICs compared with TFIID PICs (**Figure 3B**; **Figure S1G**). These results suggested that CDK7 kinase activity regulates RNAPII pause release, because TBP PICs were refractory to CDK7 inhibition, and TBP PICs will not undergo promoter-proximal pausing.^26^ To test further, we completed pausing assays in which TBP- or TFIID-containing PICs were compared ±CDK7 inhibition. As expected, RNAPII pausing was lost in TBP-containing PICs (compare 5min data in **Figure 2C** with **Figure 3C**) and the increased pausing caused by CDK7 inhibition was observed only with TFIID-containing PICs (**Figure 3D**; **Figure S1H**). These results suggest that CDK7 kinase activity enhances RNAPII pause release in a TFIID-dependent manner.

Prior biochemical studies showed functional cooperativity between Mediator and TFIID^20,32^ and reported repressive roles for TFIID ^33^ or Mediator^34^ under basal conditions (i.e. no TF activation). Although these prior studies used different experimental conditions, their results are consistent with the data summarized in **Figure 3A-D**, which reveal independent but cooperative roles for Mediator and TFIID in CDK7-dependent activation of RNAPII re-initiation. Consistent with independent roles, removal of both factors (i.e. PICs lacking Mediator, and TBP instead of TFIID) impacted CDK7-dependent runoff transcription in an additive manner, compared with loss of either factor alone (**Figure S1I, J**). Collectively, these results reveal that Mediator and TFIID cooperate to restrict and/or slow promoter-proximal RNAPII transcription, and this restriction is alleviated by CDK7 kinase activity.

The mechanistic findings from the reconstituted transcription assays are summarized as follows. 1) Overall RNAPII transcriptional output was reduced with CDK7 inhibition. 2) The reduced transcription occurred, at least in part, through reduced re-initiation in CDK7-inhibited conditions. Thus, 3) CDK7 activity promotes RNAPII re-initiation (**Figure 2**). We showed that 4) CDK7 inhibition slowed RNAPII promoter escape and 5) increased RNAPII pausing. These events (4, 5) will independently suppress RNAPII re-initiation because each event (promoter escape and pause release) will favor re-initiation by eliminating steric interference from a promoter-associated RNAPII complex. We also demonstrated that 6) the CDK7-dependent effects on RNAPII pausing required TFIID, whereas 7) Mediator contributed to the CDK7-dependent effects on promoter escape (**Figure 3**). Finally, we observed that 8) the effects of Mediator and TFIID on CDK7-dependent RNAPII transcription were additive (**Figure S1J**), consistent with the fact that promoter escape and pause release are independent regulatory events that separately contribute to RNAPII re-initiation by clearing the promoter region for additional RNAPII enzymes.

### CDK7 inhibition globally reduces RNAPII transcriptional output in human cells

Although a defined, reconstituted transcription system has advantages compared with cell-based assays, our *in vitro* assays do not match the complexity of regulatory inputs at active promoters in human cells. Therefore, we sought to further test the *in vitro* mechanistic findings in human cells using SY-5609 to selectively inhibit CDK7. Although SY-5609 has a selectivity profile as good or better than any available CDK7 inhibitor,^21,35^ we completed RNA-seq experiments in a CDK7 analog-sensitive (CDK7AS) OV90 cell line to compare with SY-5609-treated OV90 cells. As expected, the data showed highly similar effects on global gene expression (**Figure S2**), confirming the selectivity of SY-5609.

To best assess the primary impacts of CDK7 on transcription, we performed precision run-on sequencing (PRO-seq), which measures nascent transcription,^36^ with and without SY-5609. We completed the experiments in the context of IFNγ stimulation to allow evaluation of inducible and constitutive transcription. HCT116 cells were pre-treated with SY-5609 (50nM; or DMSO control) for 30min prior to a 45min IFNγ treatment. The 45min timeframe was chosen based upon prior experiments in mouse and human cells.^37^ We confirmed that SY-5609 inhibited CDK7 under these conditions (**Figure S3**), in agreement with prior results.^21^ At longer timepoints (6h), cellular ser5-P levels recovered after SY-5609 treatment, consistent with data from HAP1 cells treated with the covalent CDK7 inhibitor YKL-5-124.^14^ This recovery may reflect compensatory responses to CDK7 inhibition, but the underlying mechanisms remain unclear.

Because CDK7 inhibition was expected to globally reduce RNAPII transcription, we employed several methods to normalize the PRO-seq data, including spike-ins with *Drosophila* S2 cell nuclei (see Methods). In agreement with a recent study using TT-seq in a CDK7AS cell line,^16^ data normalization revealed that CDK7 inhibition with SY-5609 reduced RNAPII transcription by approximately 22% on average, genome-wide (**Figure 4A**), although some genes were affected more than others (see below). This result was consistent with the *in vitro* data ±SY-5609, suggesting that general defects in RNAPII promoter-proximal transcription contribute to this effect. We emphasize, however, that many additional factors, beyond PIC components, will converge on RNAPII at gene 5’-ends in cells.^1,38^

**Figure 4.**
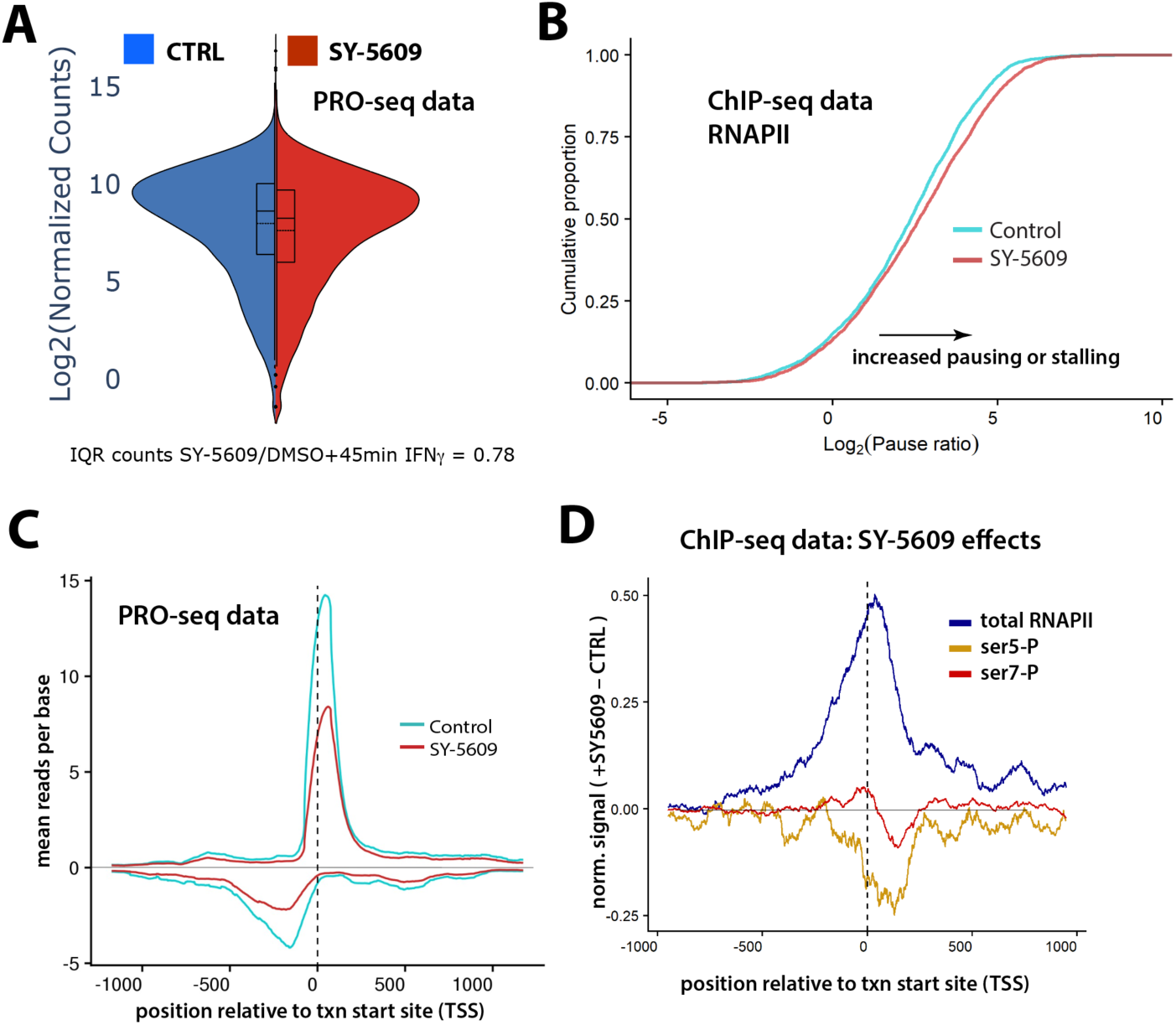
CDK7 inhibition reduces RNAPII transcription genome-wide, including at promoter-proximal regions, but increases promoter-proximal RNAPII occupancy. (A) Violin plots of the PRO-seq normalized count data. The interquartile range (IQR) counts are plotted (i.e. the middle 50% of data), showing a reduction in SY-5609-treated cells. (B) CDK7 inhibition increases RNAPII pause index (n=2400 genes; p = 1.26E^–11^). (C) Metagene plot of normalized PRO-seq reads^57^ at highly expressed genes (n=453), focused on promoter-proximal region ±SY-5609. (D) Metagene analysis of ChIP-seq data (normalized to read depth), focused on promoter-proximal region ±SY-5609 at high-occupancy RNAPII genes (n=185). Elevated RNAPII occupancy +SY-5609 coupled with reduced PRO-seq reads +SY-5609 (panel C) suggests increased levels of inactive RNAPII around the TSS. As expected, levels of RNAPII CTD ser5-P and ser7-P decrease at the TSS in SY-5609-treated cells.

After PRO-seq data normalization, several hundred transcripts were differentially expressed, with most showing reduced levels in CDK7-inhibited cells (vs. DMSO controls; **Figure S4A-C**). Interestingly, IFNγ response pathways were not down-regulated in CDK7-inhibited cells based upon GSEA (**Figure S4D**), and this was further supported by analysis at individual genes (**Figure S4C**). Instead, the down-regulated genes represented constitutively active pathways in proliferating cells (e.g. MYC targets, E2F targets, G2M targets; **Figure S4D**). These results are consistent with other studies with CDK7 inhibitors, in which stimulus-induced transcription was not negatively affected.^39,40^

### Promoter-proximal RNAPII function is disrupted in CDK7-inhibited cells

The global, genome-wide reduction in RNAPII transcription in CDK7-inhibited cells (**Figure 4A**) suggested mechanistic parallels with our *in vitro* results. To explore further, we completed a series of ChIP-seq experiments in control vs. CDK7-inhibited HCT116 cells. To match the PRO-seq experiments, cells were pre-treated with SY-5609 for 30min (50nM, vs. DMSO controls) and then stimulated with IFNγ for 45min. ChIP-seq analysis of RNAPII showed evidence of increased promoter-proximal pausing in CDK7-inhibited cells (**Figure 4B**) consistent with prior results with 1h treatment with the CDK7 inhibitor THZ1.^8^ Interestingly, increased promoter-proximal pausing was also observed at IFNγ-responsive genes (**Figure S5A**), suggesting that a common factor(s) was targeted by CDK7 at IFN-inducible genes, such as the RNAPII CTD and/or other PIC-associated factors. Our ChIP-seq results are similar to a recently published RNAPII MNase-seq experiment, which showed increased levels of PIC-associated RNAPII in CDK7-inhibited cells.^16^ A distinction is that the MNase-seq data could reliably establish RNAPII retention in the PIC, upstream of the TSS,^16^ whereas our ChIP-seq data lack this resolution.

ChIP-seq and PRO-seq are complementary techniques. Whereas ChIP-seq measures RNAPII occupancy regardless of activity (e.g. inactive complexes will be detected) PRO-seq maps all active polymerases on the genome. Comparison of our PRO-seq and ChIP-seq data at gene 5’-ends revealed differences that were consistent with decreased re-initiation in CDK7-inhibited cells, in agreement with our *in vitro* results. Specifically, metagene data showed fewer PRO-seq reads at gene 5’-ends in CDK7-inhibited cells vs. controls (**Figure 4C**), whereas ChIP-seq revealed an opposite trend: elevated levels of RNAPII at gene 5’-ends (**Figure 4D**). This disconnect between PRO-seq and ChIP-seq is consistent with an increase in stalled or inactive promoter-associated RNAPII upon CDK7 inhibition, which will reduce PRO-seq reads at gene 5’-ends.

Because CDK7 is an RNAPII CTD kinase, we also used ChIP-seq to assess genome-wide changes in CTD phosphorylation ±SY-5609. CDK7 has been shown to phosphorylate ser5 and ser7 in the RNAPII CTD^17,41^ and therefore we completed ChIP-seq experiments with antibodies targeting these modifications. Because TFIIH and CDK7 bind promoters as part of the PIC, we focused on CTD phosphorylation at gene 5’-ends. As expected, ser5-P levels declined at transcription start sites in CDK7-inhibited cells, compared with DMSO controls (**Figure 4D**). This decrease in promoter-proximal ser5-P is consistent with prior ChIP-seq experiments completed in CDK7 analog-sensitive cells.^8^ Reduced promoter-proximal ser5-P levels matched quantitative western and single cell immunofluorescence results, which showed reduced levels in CDK7-inhibited cells (**Figure S3**). Similar to ser5-P, ser7-P levels decreased at transcription start sites in CDK7-inhibited cells (**Figure 4D**), and this was consistent with quantitative western data (**Figure S3**).

### CDK7 inhibition causes readthrough transcription at gene 3’-ends

A striking result from the PRO-seq experiments was widespread “readthrough” transcription at gene 3’-ends in CDK7-inhibited cells (**Figure 5A**). Metagene analyses from PRO-seq data, focusing on long genes (over 120kb in length; **Figure 5B**), suggested that readthrough transcription was common at gene 3’-ends. This was confirmed by metagene analysis of a more general set of well-expressed genes (**Figure S5B**); furthermore, RNAPII ChIP-seq data showed elevated levels at gene 3’-ends in CDK7-inhibited cells (**Figure S5C**). Although a recent study using TT-seq and mNET-seq in CDK7AS cells concluded that CDK7 inhibition had minimal 3’-end effects,^16^ our analysis of the same data showed evidence for 3’-end readthrough transcription, similar to our results with SY-5609 (**Figure S5D, E**). The different conclusions reflect the fact that reads beyond 3’-end poly-adenylation sites were not examined previously. RNAPII 3’-end readthrough has been reported before in CDK7-inhibited cells;^8,17,18^ however, a mechanistic basis has been lacking.

**Figure 5.**
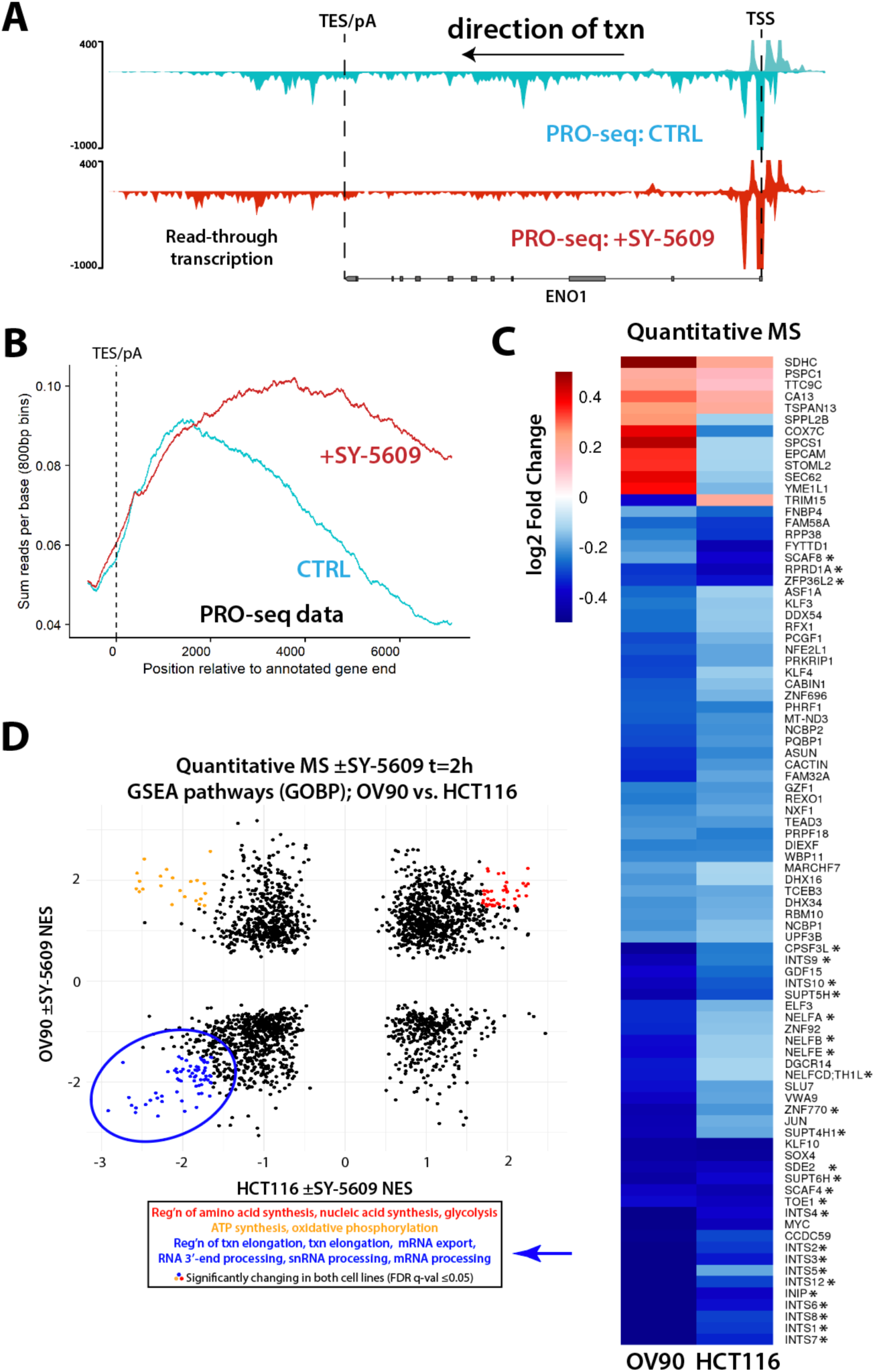
Readthrough defects and depletion of RNAPII elongation and termination factors in CDK7-inhibited cells. (A) Representative PRO-seq gene traces demonstrating increased 3’-end readthrough transcription +SY-5609. (B) Metagene analysis at long genes (>120kb, n=684) shows extensive 3’-end readthrough transcription with SY-5609 treatment. Position 0 indicates the polyA site. (C) Heatmap summarizing quantitative TMT-MS data from OV90 and HCT116 cells treated with 50nM SY-5609 for 120min (vs. DMSO controls). Proteins shown were identified as significantly changing in both OV90 and HCT116 cells (p<0.01). Asterisk: RNAPII elongation/termination factors. (D) GSEA results (GOBP set), based on TMT-MS data ±SY-5609, plotted by normalized enrichment score (NES) for OV90 and HCT116 cells. Significantly upregulated or downregulated pathways (FDR q < 0.05) are highlighted in color, and a representative subset of pathways are listed.

### Rapid loss of elongation, termination, and 3’-end processing factors upon CDK7 inhibition

Phospho-proteomics experiments in human HL60 cells identified numerous RNAPII elongation, termination, and 3’-end processing factors as high-confidence CDK7 targets.^15^ To expand upon these results, we completed quantitative mass spectrometry experiments (LC-MS/MS; TMT labeling with 4 biological replicates/condition) from two different cell lines (HCT116 and OV90). For each line, cells were treated with DMSO or SY-5609 for 2 hours under normal growth conditions (i.e. no IFNγ stimulation). Nuclei were then isolated for subsequent MS analysis. We chose a 2-hour treatment to allow time for CDK7-dependent effects to occur; MS data following 30-minute SY-5609 treatment showed few significant proteome changes (**Table S2, S3**).

Over 8000 proteins were identified in each experiment (**Table S2, S3**) and the data from HCT116 and OV90 cells yielded similar but not identical results. As shown in **Figure 5C**, a set of about 100 proteins showed reduced abundance upon CDK7 inhibition in both cell lines. Note that the magnitude of changes reported (ca. 20-50% loss) are likely minimum values given the ratio compression that occurs with TMT labeling.^42^ A pathway analysis (GSEA) of depleted proteins showed an enrichment for factors involved in RNAPII elongation, termination, and 3’-end RNA processing in each cell line (**Figure 5D**).

To build upon these results, we completed similar quantitative proteomics experiments in a CDK7AS OV90 cell line (**Table S4**); in this case, we treated cells with the ATP analog 3MB-PP1 for 4h to obtain data at a longer timepoint. The results showed a high degree of overlap with OV90 cells treated with SY-5609 (**Figure 6A, B**), revealing that elongation and termination factor loss persists at least 4 hours with CDK7 inhibition. The similar results from the 3MB-PP1 analog in CDK7AS cells vs. SY-5609 treatment also underscores SY-5609 selectivity for CDK7.^21^

**Figure 6.**
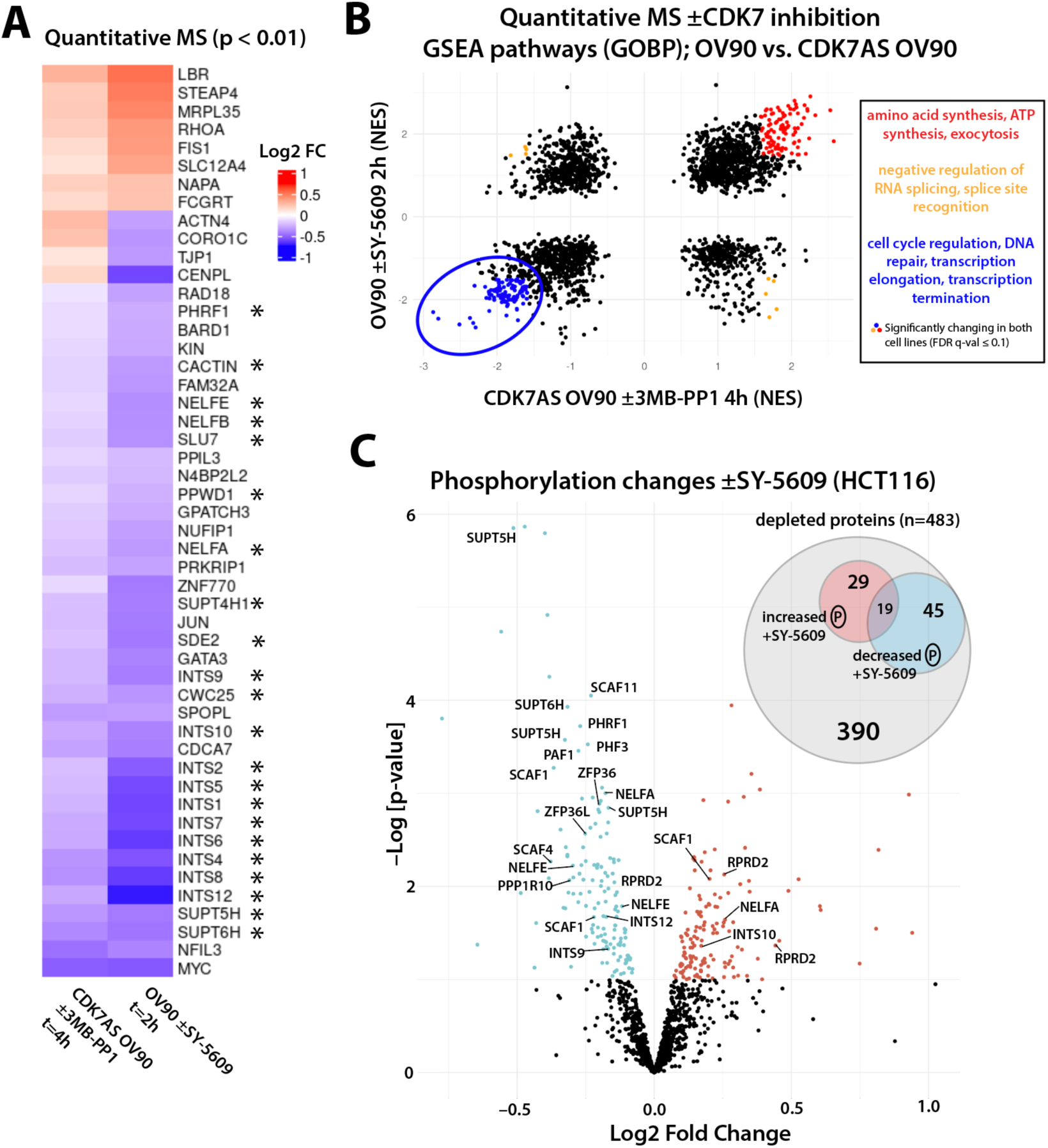
CDK7 inhibition (CDK7AS line) depletes RNAPII elongation and termination/3’-end processing factors, many of which are phosphorylated by CDK7. (A) Heatmap showing significantly changing (p < 0.01) protein abundance from quantiative TMT-MS experiments in CDK7AS OV90 cells ±3MB-PP1 (10µM, t=4h) or OV90 cells ±SY-5609 (50nM, t=2h). Proteins associated with transcription elongation, mRNA splicing, and termination are starred. (B) GSEA results (GOBP set), based on TMT-MS data ±SY-5609 (y-axis; OV90) or ±3MB-PP1 (x-axis; CDK7AS OV90), plotted by their normalized enrichment score (NES). Significantly upregulated or downregulated pathways (FDR q < 0.05) are highlighted in color, and a subset of pathways are listed at right. (C) Volcano plot showing proteins with phosphorylation changes ±SY-5609 (50nM, 1h) in HCT116 cells. For simplicity, only proteins that were also nuclear depleted +SY-5609 are included here. Among all proteins depleted with statistical confidence, about 20% showed phospho-site changes (93/483; Venn diagram inset).

We confirmed the depletion of a subset of factors (INTS2A, SPT5, and PPP1R10/PNUTS) with quantitative western blots (**Figure S6A**). To begin to assess whether factor depletion resulted from proteasomal degradation, we measured the abundance of four proteins in cells treated with the proteasome inhibitor MG132. As shown in **Figure S6B**, KLF10 was stabilized by proteasome inhibition whereas INTS2A, SPT5, and PNUTS were not affected. We obtained all quantitative proteomics data from nuclear extracts; we did not examine cytoplasmic fractions. Thus, depletion of factors upon CDK7 inhibition might reflect nuclear export rather than proteolytic degradation. To address this question, we completed quantitative westerns with cytoplasmic fractions, ±SY-5609 (t=2h). The data showed elevated levels of KLF10, SPT5, and PNUTS (**Figure S6C, D**), suggesting these proteins are exported to the cytoplasm upon CDK7 inhibition. Although the western data represent only a fraction of factors whose nuclear levels changed upon CDK7 inhibition, our results suggest that CDK7 regulates the nuclear abundance of RNAPII regulatory proteins via nuclear export and/or proteasomal degradation.

### Elongation & termination factor depletion causes 3’-readthrough defects

We next asked whether targeted depletion of a specific factor would mimic the 3’-end readthrough observed in CDK7-inhibited cells. We found that this question had already been demonstrated in many instances. For example, readthrough transcription at protein-coding gene 3’-ends occurred upon depletion of SPT5,^43^ the PAF complex,^43^ PNUTS/PPP1R10,^44,45^ SCAF4,^46^ NELF,^47^ or the Integrator complex^48^ in human cells. Thus, depleted levels of RNAPII elongation and termination factors provides a mechanistic basis for the readthrough defects in CDK7-inhibited cells. This mechanism is further supported by the fact that readthrough defects are amplified at long genes (compare **Figure 5B** with **Figure S5B**), which require more time for RNAPII to reach the 3’-end, and therefore more time for factor depletion to occur. Although no single factor was completely depleted in CDK7-inhibited cells, we hypothesize that reduced nuclear levels of many factors at once contributes to the 3’-readthrough defects. Finally, we note that changes in RNAPII CTD phosphorylation may also contribute to readthrough defects upon CDK7 inhibition.

### Depleted elongation & termination factors are phosphorylated by CDK7

Finally, we completed quantitative phospho-proteomics experiments (LC-MS/MS; TMT labeling with 5 biological replicates/condition) in the same cell lines (HCT116 and OV90; **Table S5, S6**). Cells were treated with DMSO or SY-5609 for 1 hour under normal growth conditions. The shorter 1h timepoint was chosen to better identify direct vs. indirect effects, while allowing time for CDK7-dependent changes to occur. As shown in **Figure 6C**, many factors that showed nuclear depletion in CDK7-inhibited cells are high-confidence CDK7 targets, and this was observed in OV90 cells as well (**Figure S7**). This result suggests a direct role for CDK7 in controlling the nuclear abundance of RNAPII regulatory factors.

## DISCUSSION

The results described here reveal mechanistic insights about how CDK7 regulates RNAPII function at gene 5’-ends and 3’-ends, through a combination of biochemical and cell-based experiments (**Figure 7**). Biochemical reconstitution of transcription initiation showed that RNAPII was “slow out of the gate” with CDK7 inhibition, and this was sensitive to PIC factors Mediator and TFIID. The CDK7-dependent effects on promoter escape are especially informative, as this regulatory intermediate cannot be measured with sequencing methods because the reads are too short to reliably map to the genome. That said, our ChIP-seq and recently published MNase-seq^16^ data show elevated RNAPII levels at the promoter upon CDK7 inhibition, coincident with reduced promoter-proximal transcription, as shown here with PRO-seq (**Figure 4C**). Because PRO-seq experiments use sarkosyl, promoter-proximal paused RNAPII is released, apparently through disruption of protein-protein interactions that stabilize RNAPII pausing. Accordingly, PRO-seq yields elevated reads at gene 5’-ends under normal conditions because paused RNAPII complexes can resume transcription.^49^ This was not observed in CDK7-inhibited cells (**Figure 4C**), suggesting that a subset of promoter-associated RNAPII is not paused, but instead exists in a distinct, inactivated state. An inactive PIC-associated RNAPII (i.e. upstream of the TSS) described by Velychko et al.^16^ can explain this discrepancy, but other possibilities include transcriptionally engaged backtracked RNAPII enzymes, which could similarly block re-initiation. We hypothesize that slowed or stalled promoter-proximal RNAPII transcription, observed with CDK7 inhibition in our reconstituted system, may contribute to this effect, but further study is needed. Reduced RNAPII initiation and re-initiation appears to be the primary means by which global RNAPII transcription is rapidly attenuated in CDK7-inhibited cells.

**Figure 7.**
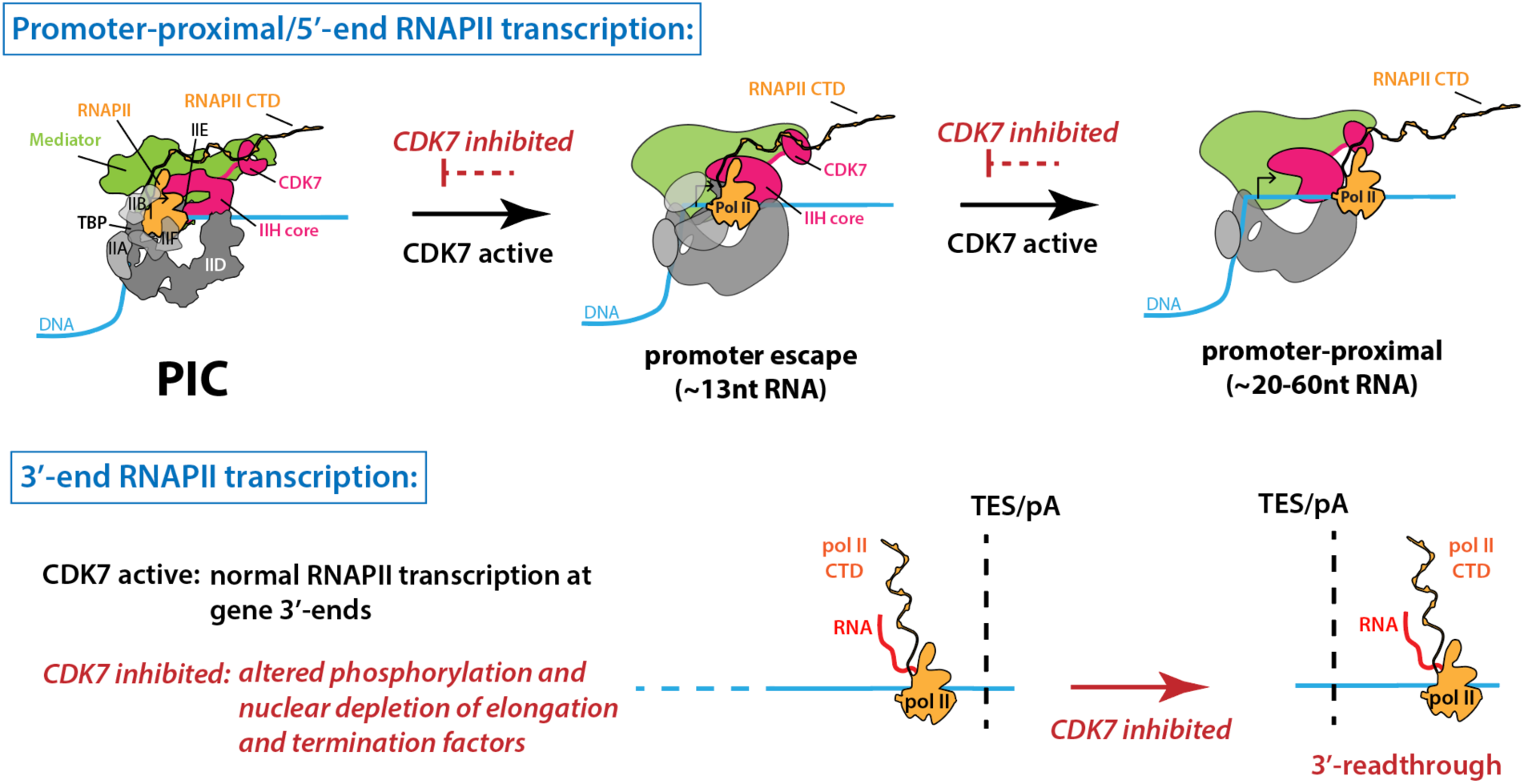
Model: CDK7 regulates RNAPII transcription at gene 5’-ends and 3’-ends through diverse mechanisms. At 5’-ends, CDK7 inhibition slows or stalls RNAPII in the promoter escape and promoter-proximal pause regions; this prevents re-initiation and may contribute to RNAPII inactivation at promoters in CDK7-inhibited cells.^16^ Separately, CDK7 inhibition causes nuclear depletion of RNAPII elongation and termination factors, which is sufficient to trigger 3’-readthrough transcription. CDK7 inhibition also causes phospho-site changes in many elongation and termination factors, which may independently contribute to the 3’-readthrough defects. PIC factors are represented as globular in the promoter escape and pause regions based upon cryo-EM data that indicate structural dynamics post-initiation.^4,50,51^

We hypothesize that the CDK7-dependent effects on promoter-proximal transcription are mediated primarily through the RNAPII CTD, because i) the CTD is a major target for the CDK7 kinase, ii) CTD phosphorylation disrupts Mediator-RNAPII interactions^8,10^ and iii) Mediator, TFIID, and RNAPII coordinate PIC assembly.^2-4^ However, TFIID and Mediator subunits are also phosphorylated by CDK7, as are other PIC factors TFIIB, TFIIE and TFIIF.^15,18^ Major structural reorganization occurs during RNAPII initiation and promoter escape, and we speculate that CDK7-dependent phosphorylation of PIC factors may be required to help coordinate structural transitions. Although cryo-EM data could build upon the mechanistic insights described here, this may be intractable given that Mediator, TFIID, and TFIIH undergo large-scale structural re-arrangements and do not adopt ordered structures during RNAPII initiation and promoter escape.^4,50,51^

Despite decades of research, our understanding of how CDK7 regulates RNAPII transcription remains incomplete. Here, through quantitative proteomics in 3 different cell lines, using structurally distinct inhibitors, we discovered that CDK7 controls the nuclear abundance of a broad spectrum of RNAPII elongation and termination factors (**Figure 5C, D**; **Figure 6A, B**), unveiling a completely new means by which CDK7 controls RNAPII function. Phosphoproteomics data showed that many of these factors were high-confidence targets of CDK7 (**Figure 6C**), providing additional mechanistic insights. For example, direct phosphorylation by CDK7 suggests that the activity of elongation and/or termination factors may be regulated by CDK7, in addition to their nuclear abundance.

Why are RNAPII elongation and termination factors depleted in the nucleus upon CDK7 inhibition? Disruption of CDK7 activity will negatively impact processing of RNAPII transcripts. Work from our lab and others has shown that active CDK7 ensures reliable 5’-end capping, splicing, and 3’-end formation.^8,13,15,17,18,52,53^ Defects in any of these processes would yield a transcriptome that is unstable and potentially pathogenic. Although the mechanisms by which CDK7 controls nuclear abundance of RNAPII regulatory factors are likely complex and protein-specific, a preliminary analysis suggests that factor depletion occurs via degradation or nuclear export (**Figure S6**). Nuclear depletion of RNAPII regulatory factors may serve to broadly reduce global transcription until CDK7 activity is restored.

It has been hypothesized that 3’-end readthrough transcription is a common cellular response to diverse types of stress,^54^ and that 3’-readthrough transcripts may be retained in the nucleus and later spliced and processed post-transcriptionally, to accelerate a return to homeostasis after stress.^55^ It is remarkable that the “master regulators” of RNAPII elongation, termination, and 3’-end processing (e.g. DSIF, SPT6, PAF complex, PNUTS, SCAF8, Integrator complex) rely on CDK7 activity to maintain nuclear localization and/or cellular stability. We speculate that regulated suppression of CDK7 activity represents a common mechanism by which cells establish 3’-end readthrough transcription during diverse stress responses.

### Limitations of this study

Although biochemical reconstitution of RNAPII transcription is helpful for assessing CDK7-dependent regulatory functions, the PIC factors used here do not recapitulate the complexity of proteins, protein complexes, nucleic acids, and metabolites that can influence gene expression in human cells. Additional transcription-associated kinases and other factors regulate RNAPII function in cells, but we only examine the PIC factors here. An improved mechanistic understanding resulted from combining *in vitro* experiments with cell-based assays, but only a few cell lines were used here, and only under specific contexts. Moreover, our transcriptomics data were obtained after 75min CDK7 inhibition, which will miss CDK7-dependent effects across longer timeframes.

## RESOURCE AVAILABILITY

### Lead contact

Further information and requests should be directed to and will be fulfilled by the lead contact, Dylan Taatjes.

### Material availability

Plasmids or reagents generated in this study are available from the lead contact upon request.

### Data and code availability

PRO-seq, ChIP-seq and RNA-seq data have been deposited at Gene Expression Omnibus (GEO) as GSE228346, GSE262536, GSE261575, GSE261649, and GSE268531. All original code has been deposited at https://github.com/Dowell-Lab/CDK7_inhibition This paper analyzes existing, publicly available data, accessible at Short Read Archive (Parent GEO accession GSE218269, TT-seq SRA sample accessions SRR22327405-SRR22327416; NET-seq SRA sample accessions SRR22327385-SRR22327391).

## Acknowledgments

We thank David Bentley (UC Anschutz School of Medicine) for helpful comments on the manuscript, Jason Marineau (Syros Pharmaceuticals) for project support and Theresa Nahreini (UC-Boulder) for cell culture assistance. We thank Jim Goodrich (UC-Boulder) for providing TAF4 antibodies for TFIID purification, and R. Tjian (UC-Berkeley) for ERCC3 antibodies for TFIIH purification. This work was supported by the NIH (R35 GM139550 to DJT; R01 GM148613 to JFK) and the NCI (F31 CA254478A to OL). Some funding support was provided by Syros Pharmaceuticals to DJT. We acknowledge funding support for instrumentation (S10OD025267).

## Author contributions

Experimental design: OL, LS, JR, TJ, CCE, MN, JFK, RDD, DJT

Data collection and analysis: OL, LS, JR, TJ, JKR, MN, GSS, KC, FB, CCE, AH, MBS, LPW

Supervision and funding: SS, RDD, DJT

Writing: OL, DJT

## Supplemental information

Document S1. Figures S1-S7

**Table S1.** MS data from immobilized template + nuclear extract experiments.

**Table S2.** Quantitative MS data for HCT116 ±SY-5609; t=30min & t=120min.

**Table S3.** Quantitative MS data for OV90 ±SY-5609; t=30min & t=120min.

**Table S4.** Quantitative MS data for CDK7AS OV90 ±3MB-PP1; t=4h.

**Table S5.** Quantitative phospho-proteomics MS data for HCT116 ±SY-5609; t=60min.

**Table S6.** Quantitative phospho-proteomics MS data for OV90 ±SY-5609; t=60min.

## STAR METHODS

### KEY RESOURCES TABLE

| Antibodies |  |  |  |
| --- | --- | --- | --- |
| Target | Manufacturer | Catalog # |  |
| RNAPII/RPB1 | Cell Signaling Technology | 14958 |  |
| Mediator/MED23 | Bethyl | 50-155-7590 |  |
| TFIIH/XPB | Cell Signaling Technology | 8746 |  |
| TFIIF/GTF2F1 | Cell Signaling Technology | 8080 |  |
| TFIIE/GTF2E1 | proteintech | 24405-1-AP |  |
| TFIIA/GTF2A1 | Thermo Fisher Scientific | PA5-102739 |  |
| TFIID/TBP | proteintech | 22006-1-AP |  |
| TFIIB/GTF2B | Cell Signaling Technology | 4169 |  |
| SPT5 | Santa Cruz Biotechnology | Sc-28678 |  |
| INTS2A | Santa Cruz Biotechnology | Sc-514945 |  |
| KLF10 | ProteinTech | 29709-1-AP |  |
| PPP1R10 | Invitrogen | PA5-48110 |  |
| phospho-CTD ser-5 | Millipore Sigma | 04-1572 |  |
| phospho-CTD ser-7 | Millipore Sigma | 04-1570 |  |
| phospho-CTD ser-2 | Millipore Sigma | 04-1571 |  |
| RNA Polymerase II Subunit B2 | GeneTex | GTX102535 |  |
| goat anti-rabbit (Alexa Flour 488) | Invitrogen | A78953 |  |
| goat anti-rat (Alexa Fluor 647) | Invitrogen | A21247 |  |
| Chemicals, Peptides, and Recombinant Proteins |  |  |  |
| Name | Manufacturer | Catalog # |  |
| flavopiridol | Sigma-Aldrich | F3055 |  |
| SY-5609 | Syros Pharmaceuticals | proprietary |  |
| SY-351 | Syros Pharmaceuticals | proprietary |  |
| Lambda Protein Phosphatase | New England Biolabs | P0753 |  |
| Lys-C/Trypsin | Promega | V5071 |  |
| GlycoBlue | Invitrogen | AM9515 |  |
| α32PCTP | Revvity | BLU008H001MC | |
| γ32P-ATP | Revvity | BLU502A100UC | |
| CAK (CDK7/CCNH/MAT1) | Millipore | 14-476M |  |
| IFNγ | Life Technologies | PHC4031 | |
| Benzamidine | Sigma | B6506-100G |  |
| Sodium Metabisulfite | Sigma | 255556-100G |  |
| Phenylmethylsulfonyl Fluoride | American Bioanalytical | AB01620 |  |
| Aprotinin | Sigma-Aldrich | A6279-10ML |  |
| Trizol reagent | Invitrogen | 15596018 |  |
| RIPA lysis buffer | Thermo Scientific | 89900 |  |
| Benzonase | Millipore Sigma | 101697 |  |
| Cell Lines |  |  |  |
| Lineage | Manufacturer | Catalog # | Sex |
| OV90 | ATCC | CRL-3585 | female |
| OV90 CDK7AS | Syros Pharmaceuticals |  | female |
| HCT116 | ATCC | CCL-247 | male |
| Other |  |  |  |

| Name | Manufacturer | Catalog # |
| --- | --- | --- |
| DNA Gel Extraction Kit | Omega Bio-Tek | D2500 |
| PCR clean-up kit | Omega Bio-Tek | D6492 |
| RNeasy kit | Qiagen | 74104 |
| KAPA mRNA HyperPrep Kit | Roche | 50-196-5308 |
| M-280 Dynabeads | Thermo Fisher Scientific | 11205D |
| AMPure XP beads | Beckman | A63881 |
| carboxylate-functionalized speedbeads | Cytiva Life Sciences | 29729998 |
| 4%–20% gradient gel | BioRad | 4561096 |
| Revert 700 Total Protein Stain | Li-Cor | 926-11021 |
| **Software** |  |  |
| Name | Version | Application |
| Cytiva ImageQuant | v10.2 | Gel quantitation |
| RNAseq-Flow |  | RNA-seq/PRO-seq/ChIP-seq |
| fastQC | v0.11.8 | RNA-seq/PRO-seq/ChIP-seq |
| preseq | v2.0.3 | RNA-seq/PRO-seq/ChIP-seq |
| samtools | v1.8 | RNA-seq/PRO-seq/ChIP-seq |
| bbduk | v38.05 | RNA-seq/PRO-seq/ChIP-seq |
| Hisat2 | v2.1.0 | RNA-seq/PRO-seq/ChIP-seq |
| DESeq2 | v3.6.0 | RNA-seq/PRO-seq/ChIP-seq |
| NascentFlow | v1.3 | PRO-seq/ChIP-seq |
| nextflow | 21.02.0.edge | PRO-seq/ChIP-seq |
| sratools | 2.8.0 | PRO-seq/ChIP-seq |
| preseq | 2.0.3 | PRO-seq/ChIP-seq |
| java | 1.8.0_292 | PRO-seq/ChIP-seq |
| picard_gc | 2.6.0 | PRO-seq/ChIP-seq |
| picard_dups | 2.6.0 | PRO-seq/ChIP-seq |
| bedtools | 2.28.0 | PRO-seq/ChIP-seq |
| igvtools | 2.3.75 | PRO-seq/ChIP-seq |
| mpich | 3.2.1 | PRO-seq/ChIP-seq |
| gcc | 7.1.0 | PRO-seq/ChIP-seq |
| python | 3.6.8 | PRO-seq/ChIP-seq |
| numpy | 1.19.5 | PRO-seq/ChIP-seq |
| GSEA | v4.3.2 | mass spec |
| R | 4.2.3 |  |

### METHOD DETAILS

#### Immobilized template assays

The native HSPA1B template comprised 500bp upstream of the transcription start site (TSS) and 127bp of downstream sequence. The template was biotinylated at the 5’ end using a biotinylated forward primer during PCR amplification of a plasmid containing the native HSP70 sequence. The template DNA was purified using DNA gel extraction (Omega Bio-Tek, cat. #D2500) and PCR clean-up kits (Omega Bio-Tek, cat. D6492) to ensure ultra-pure DNA. Template immobilization on M-280 Dynabeads (Thermo Fisher Scientific, cat. #11205D) was performed following the manufacturer’s instructions. The immobilized template was quantitated via NanoDrop. Each assay contained 1µg immobilized template DNA (IMT) in a total volume of 100 µL. Purified HSF1 was incubated with the IMT in activator buffer (20 mM HEPES, 10% glycerol, 70 mM KCl, 8 mM MgCl_2_, 0.1 mM EDTA, 0.25µg/µL BSA, 0.0125% NP-40, 0.5 mM DTT, 0.3µM ZnCl_2_) at room temperature for 30 minutes on a HuLa Mixer (Thermo Fisher Scientific, cat. #15920D) to ensure constant bead agitation. The IMT was then captured on a magnetic strip, and the supernatant was discarded. The IMT was immediately resuspended in 100µL wash buffer (20 mM HEPES, 10% glycerol, 70 mM KCl, 8 mM MgCl_2_, 0.1 mM EDTA, 0.25µg/µL BSA, 0.0125% NP-40, 0.5 mM DTT) by gentle vortexing, followed by a quick spin on a microcentrifuge. After reapplying the magnetic strip and removing the supernatant, the IMT was incubated with 100µL PIC buffer containing HeLa NE (13.2 mM HEPES, 13.2% glycerol, 66 mM KCl, 8 mM MgCl_2_, 0.132 mM EDTA, 0.25µg/µL BSA, 0.0125% NP-40, 0.5 mM DTT, 400ng of sheared salmon sperm DNA). Each assay contained 630μg of HeLa NE; nuclear extracts were generated as described.^58^ This mixture was incubated at room temperature for 60 minutes on a HuLa Mixer, with resuspension every 20 minutes to prevent bead settling. After a quick spin, the IMT was placed on a magnetic strip to remove the supernatant. The IMT was immediately washed with 100µL wash buffer and resuspended in 100µL transcription buffer (20 mM HEPES, 10% glycerol, 70 mM KCl, 8 mM MgCl_2_, 0.1 mM EDTA, 0.5 mM DTT, 10µg of sheared salmon sperm DNA) and incubated on a thermoshaker at 30°C for 2 minutes, prior to NTP addition (NTPs added to final concentrations of 0.5mM for CTP, GTP, and UTP, and 1mM for ATP). For +inhibitor experiments, flavopiridol (500nM final concentration; Sigma-Aldrich F3055), SY-5609 (250nM) or SY-351 (100nM) were incubated for 10min prior to NTP addition. Reactions were incubated with NTPs at 30°C for 10 minutes at 1000 RPM on a thermoshaker. The IMT reactions were then immediately placed on the magnetic strip and the supernatant was removed, followed by immediate incubation with 100µL wash buffer, then resuspended in SDS loading buffer, and worked up for western blot or mass spectrometry analysis. For western blot, proteins were resolved using 4-20% SDS-PAGE gels (Bio-Rad, cat. #4561094EDU) and transferred to PVDF membranes (Thermo Fisher Scientific, cat. #88518). Membranes were blocked in 5% milk (in TBS-T) for 1h and cut into strips for overnight incubation with primary antibodies. The antibodies used were as follows: RNAPII/RPB1 (Cell Signaling Technology clone D8L4Y, cat. #14958), Mediator/MED23 (Bethyl, cat. #50-155-7590), TFIIH/XPB (Cell Signaling Technology clone 2C6, cat. #8746), TFIIF/GTF2F1 (Cell Signaling Technology, cat. #8080), TFIIE/GTF2E1 (proteintech, cat. #24405-1-AP), TFIIA/GTF2A1 (Thermo Fisher Scientific, cat. #PA5-102739), TFIID/TBP (proteintech, cat. #22006-1-AP), TFIIB/GTF2B (Cell Signaling Technology clone 2F6A3H4, cat. #4169). SuperSignal West Pico PLUS Chemiluminescent Substrate (Thermo Fisher Scientific, cat. #34579) was used for protein visualization following 1h secondary antibody incubation and TBS-T washes.

##### Phosphatase experiment

After NTP addition and incubation (described above), IMT samples were placed on a magnetic strip, washed with 100µL wash buffer, and resuspended in 5µL 10× NEBuffer for Protein MetalloPhosphatases, 5µL 10 mM MnCl_2_, 35µL water, and 5µL Lambda Protein Phosphatase (New England Biolabs, cat. #P0753). Samples were incubated at 30°C for 10 minutes at 1000 RPM on a thermoshaker. Following incubation, the samples were placed on a magnetic strip, the supernatant removed and subsequently resuspended in SDS loading buffer for western blot analysis.

#### Sample Preparation for Immobilized Template MS Proteomics

Proteins bound to immobilized templates were denatured, reduced and alkylated using 5% (w/v) sodium dodecyl sulfate (SDS), 10 mM tris(2-carboxyethylphosphine) (TCEP), 40 mM 2-chloroacetamide, 50 mM Tris-HCl, pH 8.5 with boiling 10 minutes, then incubated shaking at 2000 rpm at 37°C for 30 minutes. Proteins were digested using the SP3 method.^59^ Briefly, 1000 µg carboxylate-functionalized speedbeads (Cytiva Life Sciences) were added followed by the addition of acetonitrile to 80% (v/v) inducing binding to the beads. The beads were washed twice with 80% (v/v) ethanol and twice with 100% acetonitrile. Proteins were digested in 50 mM Tris-HCl, pH 8.5, with 1.0 µg Lys-C/Trypsin (Promega) and incubated at 37°C overnight. Tryptic peptides were desalted with the addition of 95% (v/v) acetonitrile binding the peptides back to the beads and washed once with 100% acetonitrile. Peptides were collected from the beads with two elutions of 1% (v/v) trifluoroacetic acid, 3% (v/v) acetonitrile. Cleaned-up peptide were then dried in a speedvac vacuum centrifuge and stored at -20°C until analysis. Tryptic peptides were suspended in 3% (v/v) ACN, 0.1% (v/v) trifluoroacetic acid (TFA) and directly injected onto a reversed-phase C18 1.7 µm, 130 Å, 75 mm X 250 mm M-class column (Waters), using an Ultimate 3000 nanoUPLC (Thermos Scientific). Peptides were eluted at 300 nL/minute with a gradient from 2% to 15% ACN in 40 minutes then to 40% ACN in 5 minutes and detected using a Q-Exactive HF-X mass spectrometer (Thermo Scientific). Precursor mass spectra (MS1) were acquired at a resolution of 120,000 from 350 to 1550 m/z with an automatic gain control (AGC) target of 3E6 and a maximum injection time of 50 milliseconds. Precursor peptide ion isolation width for MS2 fragment scans was 1.4 m/z, and the top 12 most intense ions were sequenced. All MS2 spectra were acquired at a resolution of 15,000 with higher energy collision dissociation (HCD) at 27% normalized collision energy. An AGC target of 1E5 and 100 milliseconds maximum injection time was used. Dynamic exclusion was set for 5 seconds with a mass tolerance of ±10 ppm. Rawfiles were searched against the Uniprot Human database UP000005640 downloaded 08/26/2022 using MaxQuant v. 2.0.3.0. Cysteine carbamidomethylation was considered a fixed modification, while methionine oxidation and protein N-terminal acetylation were searched as variable modifications. All peptide and protein identifications were thresholded at a 1% false discovery rate (FDR).

#### In vitro transcription assays

All incubations in the transcription assay are at 30°C. Transcription was conducted on linearized native human HSPA1B promoter extending from -504 to +216 around the annotated transcription start site, essentially as described ^26^. Briefly, the template was first incubated with 400nM HSF1 for 30 minutes in buffer containing 20mM HEPES, 80mM MgCl_2_, 280nM ZnCl_2_, and 5% glycerol. PIC factors (TFIIA, TFIIB, TFIID, TFIIE, TFIIF, TFIIH, Mediator, RNAPII) were combined into a master mix that was then added to the promoter templates and incubated for 15 minutes to allow PICs to assemble. For experiments ±SY-5609, the compound (or DMSO control) was pre-incubated with TFIIH for 15min on ice prior to combining into the master mix. Transcription was initiated by the addition of NTPs. For “single-round” experiments, sarkosyl (0.2%) was added 1min after the addition of NTPs. Transcription reactions were stopped with the addition of 150µL of Stop Buffer (20mM EDTA, 200mM NaCl, 1% SDS, 250mM ammonium acetate, GlycoBlue (Invitrogen #AM9515)). The experimental variations are described below:

##### Standard Runoff Experiments

2µL of stock NTPs (6.25mM ATP, GTP, UTP, 1.25mM CTP, 0.01mCi α32P-CTP (Revvity #BLU008H001MC)) were added to each reaction (final volume 20µL) and allowed to incubate at 30°C for various times.

##### Pausing assays

2µL of stock NTPs (6.25mM ATP, GTP, UTP, 0.01mCi α32P-CTP) were added to each reaction followed by adding 2µL of 0.1mM CTP 1 minute later (final volume 22µL). Transcription continued for an additional 4 minutes at 30°C or was immediately stopped.

#### Transcription Gel Quantitation

Band intensity was used to calculate transcript quantities based upon ^32^P-labeled RNAs. Bands were quantified using Cytiva ImageQuant software (v10.2). First, the background was subtracted with a rolling-ball radius of 2. Band intensity was measured as the area under the curve minus the background. Runoff transcripts were defined between 150-216nt, paused transcripts between 20-70nt, and promoter-associated transcripts 5-15nt. Though we acknowledge that promoter escape occurs at 12-13nt, the smallest transcript length that could be accurately determined from a marker reference was 15nt.

##### Normalization calculations

Because the *in vitro* transcription assays use ^32^P-labeled CTP, we can visualize all transcripts. However, a caveat is that ^32^P-CTP decays with a half-life of approximately 14 days; therefore the signal intensity will vary accordingly. Moreover, intensities will vary based upon exposure to the phosphor-imager screen. Completion of experiments with many biological replicates required multiple different assays completed on different days. To account for potential variability in ^32^P signal, all runoff experiments were normalized to a “base PIC” internal control that contained all PIC factors except Mediator. This “base PIC” experiment was included in at least 2 replicates for all experiments. This internal control provided a reference that allowed reactions to be accurately compared across different experiments.

For pausing assays, a pause ratio (total intensity of paused transcripts/total intensity of runoff transcripts) or promoter escape ratio (total intensity of promoter-associated transcripts/total intensity of runoff transcripts) was calculated for control and +SY-5609 reactions. No additional normalization to “base PIC” was required here because the data were calculated as a ratio, which inherently normalizes experimental variation. The regions defined for paused, runoff, and promoter-associated transcripts were 20-70nt, 150-216nt, and 5-15nt, respectively.

##### In vitro kinase assays

The kinase assay with purified CAK (Millipore CDK7/CCNH/MAT1; 14-476M) and RNAPII Rpb1 CTD was conducted as follows. Mouse GST-tagged Rpb1 CTD (full length, 52 heptad repeats), 10nM final) was incubated in buffer (0.1M KCl, 25mM Tris-HCl pH 7.9, 10mM MgCl_2_, 2mM DTT) for 10 minutes at 37°C. Meanwhile, purified CAK module (50nM final) was pre-incubated with 500nM SY-5609 (50nM final for 20µL reaction volume) for 15 minutes on ice. CAK module was then added to the GST-CTD mixture and phosphorylation was initiated by the addition of 600µM ATP and 10µCi γ32P-ATP and incubation at 37°C. For promoter-bound PIC kinase assays, PICs were assembled as described for the *in vitro* transcription assays. Next, 2µL of stock NTPs (6.25mM CTP, GTP, UTP, 0.01mCi γ32P-ATP) was added, followed by 2µL of 6mM ATP 1 minute later (final volume 22µL). Thus, phosphorylation was evaluated rather than transcription while replicating *in vitro* transcription assay conditions. Phosphorylation continued for an additional 4 minutes or 29 minutes at 37°C. Reactions were stopped at designated timepoints with the addition of 4x Laemmli Buffer and boiled for 5 minutes. Samples were then run on a 4%–20% gradient gel (BioRad 4%–20% Mini-Protean TGX gel, 15-well). Gels were dried and exposed to evaluate autorad signal. ImageJ software was used to measure autorad signal.

#### RNA-seq

##### Cell culture and treatments

OV90 (ATCC, CRL-3585) and OV90 CDK7AS cells were cultured in 1:1 MCDB 105 medium (Cell Applications, 117-500) to medium 199 (Sigma-Aldrich, M4530-500ML), supplemented with 15% fetal bovine serum and 1% Penicillin-Streptomycin and grown on 150 mm^2^ dishes. All cells were cultured at 37°C with 5% CO_2_. Cells were grown to ∼75% confluency and treated with 50nM SY-5609 or equivalent volume DMSO (0.01% DMSO) for 30min and heat shocked as described^60^ for 30 minutes. Briefly, heat shock was performed by removing half of the conditioned media, to be heated to 49°C, then adding back to flasks containing cells (at 37°C) such that the media temperature on the cells achieved 42°C instantly. Cell culture flasks (T-225) were then submerged in a 42°C water bath for 30min, followed by a 60min recovery at 37°C. Cells were then washed three times with cold PBS, lysis buffer containing SUPERaseIN (RNase inhibitor) was added, and cells harvested by scraping in cold lysis buffer, followed by RNA isolation in TRIzol.

The heat shock treatments were intended for a separate project, but we were asked to address concerns about SY-5609 selectivity for this article. The RNA-seq comparisons here address the selectivity question in a different way compared with published kinome-wide screens.^21^ We emphasize that both cell lines were treated identically, and both lines are OV90; however, the CDK7AS line underwent clonal selection after genome editing and thus can be considered a genetically distinct line. Another confirmation of SY-5609 selectivity derives from the quantitative proteomics comparisons in OV90 and CDK7AS OV90 cells (**Figure 6A, B**).

##### RNA-seq library preparation and sequencing

Upon harvesting, RNA was extracted with TRIzol reagent, according to the manufacturer’s instructions. DNaseI digestion was then performed with the RNeasy kit (Qiagen) using the provided on-column protocol.^15^ RNA was eluted in 50µL RNase-free water and samples analyzed by Qubit and TapeStation. Samples were concentrated to 300-400ng/µL, and poly(A) selection was performed with the KAPA mRNA HyperPrep Kit with 1µg of RNA containing 1:100 diluted ERCC RNA spike-in mix (Invitrogen Thermo Fisher). Samples were then sequenced on the NovaSEQ6000, using a paired-end 150bp cycle (2x150).

##### Mapping & QC

RNA-seq mapping and QC were by the Nextflow pipeline RNAseq-Flow (https://github.com/Dowell-Lab/RNAseq-Flow). QC was performed by fastQC (v0.11.8), and preseq (v2.0.3). QC and file type conversions were performed with samtools (v1.8). Adapter trimming was performed by bbduk from bbmap (v38.05). Mapping was performed by Hisat2 (v2.1.0).^61^ Mapping was performed to the human reference genome hg38 (GRCh37). Differential expression analysis was conducted with the R package DESeq2 (v3.6.0) and expression was normalized to ERCC RNA spike-in.

#### Precision Run-on Sequencing (PRO-seq)

##### Cell culture and treatments

HCT116 cells were grown in McCoy’s 5A media (10% fetal bovine serum, 1% Penicillin/streptomycin) to ∼75% confluency on 500cm^2^ plates (Corning 431110). At t = 0 minutes, 50µM SY-5609 (1000x stock or DMSO volume equivalent) was added to the conditioned media and mixed thoroughly. At t = 30 minutes, 1µg/mL IFNγ (Life Technologies PHC4031; 1000x stock) or 1x phosphate-buffered saline (PBS) was added to the conditioned media and mixed thoroughly. At t = 75 minutes, the media was aspirated from the plate and the cells were washed in ice cold 1x D-PBS three times. 40mL of ice-cold Lysis Buffer (10 mM Tris–HCl pH 7.4, 2 mM MgCl_2_, 3 mM CaCl_2_, 0.5% NP-40, 10% glycerol, 1 mM DTT, 1x Protease Inhibitors (1mM Benzamidine (Sigma B6506-100G), 1mM Sodium Metabisulfite (Sigma 255556-100G), 0.25mM Phenylmethylsulfonyl Fluoride (American Bioanalytical AB01620), and 4U/mL SUPERase-In) was then added to the plate and cells were scraped with a cell-lifter (Costar) and the resuspended cells were transferred into a conical and put on ice.

##### Nuclei isolation

Resuspended cells were allowed to swell in lysis buffer, on ice, for 0.5-1hour (while the other cells were being harvested). Cells were then pelleted in a fixed-angle rotor centrifuge at 1000xg for 15 minutes. Pelleted cells were resuspended in 10mL of Lysis Buffer with a wide-bore P1000 tip and the centrifugation step was repeated. Being mostly nuclei and cell debris, the pellet was resuspended in 500µL Freezing Buffer (50 mM Tris pH 8.3, 40% glycerol, 5 mM MgCl_2_, 0.1 mM EDTA, 4U/ml SUPERase-In) and moved to a low-retention 1.5mL tube and pelleted at 2000xg for 4 minutes. The final pellet was resuspended in Freezing Buffer, checked for purity and counted under a microscope, and brought to a concentration of 10 million nuclei/100µL then aliquoted and snap frozen.

##### Library preparation

Nuclear run-on was conducted as described^62^ with ATP, GTP, and UTP at 0.125mM; however, both CTP and biotin-CTP were used at a concentration of 0.025mM. The nuclear run-on was performed at 37°C for 3min with a gentle mix at 1.5min. RNA was extracted with Trizol reagent (Invitrogen 15596018), sheared by base hydrolysis and nascent RNA was isolated with magnetic streptavidin beads (Thermo Scientific 88816). The final library clean-up and size selection was accomplished using 1X AMPure XP beads (Beckman A63881).

#### PRO-seq data analysis

##### Sequencing and initial processing

Sequencing was performed by the Nevada Genomics Center at the University of Nevada Reno. Single-end fragment libraries (100 bp) were sequenced by Illumina NextSeq 2000, demultiplexed, and converted to fastq format by the UNR facility. Sequencing data quality was assessed with the Illumina BaseSpace program.

Initial data processing was conducted with the NascentFlow pipeline (v1.3) with –flip and – single_end flags and with the following program versions: nextflow_version: 21.02.0.edge, fastqc_version: 0.11.8, bbmap_version: 38.05, hisat2_version: 2.1.0, samtools_version: 1.8, sratools_version: 2.8.0, preseq_version: 2.0.3, java_version: 1.8.0_292, picard_gc_version: 2.6.0, picard_dups_version: 2.6.0, bedtools_version: 2.28.0, igvtools_version: 2.3.75, mpich_version: 3.2.1, gcc_version: 7.1.0, python_version: 3.6.8, numpy_version: 1.19.5. Data were aligned to the human reference genome (hg38). More details about the NascentFlow pipeline can be found here doi: 10.17605/OSF.IO/NDHJ2

Millions-mapped was used to select the isoforms expressed but ultimately a more rigorous normalization was required. We tested the exogenous spike-in of S2 nuclei but had highly variable results possibly due to the challenges with consistently delivering equal amounts of nuclei to each sample, given that S2 nuclei tend to stick together. Variability may have also been due to lack of efficiency of *Drosophila* S2 cell transcription at 37°C and/or heterogeneity of S2 nuclei populations across samples. Consequently, we used an internal “virtual spike-in” (VSI) normalization method that relies on the 3’ end of long genes.^57^ The theory is that the 3’ end of a subset of genes is too long to be impacted by the longest treatment times. In this study, the longest treatment times were 75min and assuming an average rate of 3kb/min, we used genes longer than 225kb (n=956 genes). This internal normalization method was comparable to most of the exogenous spike-in samples with a few exceptions due to the hypervariability in the S2 normalization.

Differential expression analysis was conducted with the R package DESeq2 (v3.6.0). Embedded programs were used to estimate dispersions for PCA plots. Counts were generated with the utility featurecounts. Counts at 5’-ends were excluded from the analysis to avoid skewing due to 5’ peaks, generating suitable model weights for further analysis. Size factors were calculated by virtual spike-in (VSI).^57^

##### Gene set enrichment analysis

GSEA was performed with the Broad Institute’s GSEA software (v4.3.2) using the pre-ranked module. Log(2) fold-change values were used as the rank metric for all genes and compared against the Hallmark gene sets database (h.all.v2023.2.Hs.symbols.gmt) for enrichment.

##### Metagene Analysis

Transcription start site (TSS) annotation files were generated from Refseq gene annotations and filtered to specific gene lists based on differential transcription results from DESeq2. These lists included IFN-stimulated genes (n=138), the most highly expressed genes (n=1000), and genes longer than 120kb. Gene lists were manually filtered to remove genes containing clear bidirectional transcription or same-stranded genes within 7.5kb downstream of the annotated gene end, yielding 96 IFN-stimulated genes, 453 highly expressed genes, and 684 genes > 120kb. From there, ‘bedtools slop’ (bedtools version 2.28.0) was used to append 1000 bases to each side of the TSS, creating 2001-base regions. Similarly, for 3’ metagenes, 1000 bases were appended to the 5’ end of the TES and 7500 bases to the 3’ end of the TES to create 8501-base regions. Base-specific and strand-specific coverage was calculated over these regions for each sample using the command ‘bedtools genomecov’ with the flag ‘-5’ to count only the 5’-most base. A custom python script (python version 3.6.3) was then used to normalize and sum read counts across replicates, filter out genes with low read counts as well as the genes with the highest 5% read counts, and bin counts. These filtered and binned counts were used to generate metagenes.

Plotting was done in R (version 4.2.3) with the packages ggplot2 (version 3.4.3), dplyr (version 1.1.3), reshape2 (version 1.4.4) and cowplot (version 1.1.1).

#### Quantitative Mass Spectrometry

##### Nuclear extract generation

HCT116 cells (ATCC, CCL-247) were grown to approximately 75% confluency on 15cm dishes (CellStar #628160) in McCoy’s 5A media with 10% FBS and 1% Penicillin-streptomycin. OV90 cells (ATCC, CRL-3585) were cultured to approximately 75% confluency on 15cm dishes in 1:1 MCDB 105 medium (Cell Applications, 117-500) to medium 199 (SigmaAldrich, M4530-500ML), supplemented with 15% FBS and 1% Penicillin-Streptomycin. Both cell lines were cultured at 37°C with 5% CO_2_. Either DMSO or 50µM SY5609 (1000x) was added to the conditioned media and allowed to incubate at 37°C for 120 minutes. Cells were washed 3x with cold 1x D-PBS and scraped into 3mL of cold wash buffer (200mM HEPES, 150mM NaCl). Cells were then pelleted by centrifuging at 1000xg for 5 minutes and then resuspended in 200µL lysis buffer (1% glycerol, 200mM HEPES, 150mM NaCl, 0.5% NP-40, 1mM EDTA). Cells then nutated at 4°C for 30-45 minutes and nuclei were pelleted by centrifuging at 2300xg for 5 minutes. The supernatant (cytoplasmic fraction) was reserved and the nuclei were then lysed by nutation with 50µL RIPA buffer (Pierce #89900) supplemented with universal nuclease (Pierce #88700) and protease inhibitors (1mM benzonase, 1mM sodium metabisulfite, 1mM DTT) for 45 minutes. Nuclei suspensions were then sonicated in a water bath sonicator 15s on, 30s off, repeating 3x.

##### Sample Preparation for Mass Spectrometry

Protein lysates were denatured, reduced and alkylated with 5% (w/v) SDS, 1% (w/v) sodium deoxycholate, 10 mM tris(2-carboxyethylphosphine) (TCEP), 40 mM 2-chloroacetamide, 50 mM Tris-HCl, pH 8.5, and boiled at 95°C for 10 minutes then incubated shaking at 2000 rpm at 37°C for 30 minutes. Lysates were digested using the SP3 method.^59^ Briefly, 200 µg carboxylate-functionalized speedbeads (Cytiva Life Sciences) were added to approximately 100 µg protein lysate. Addition of acetonitrile to 80% (v/v) induced binding to the beads, then the beads were washed twice with 80% (v/v) ethanol and twice with 100% acetonitrile. Proteins were digested in 50 mM Tris-HCl buffer, pH 8.5, with 1 µg Lys-C/Trypsin (Promega) and incubated at 37°C overnight. Tryptic peptides were desalted using HLB Oasis 1cc (10mg) cartridges (Waters) according to the manufacturer’s instructions and dried in a speedvac vacuum centrifuge. Approximately 7 µg of tryptic peptide from each sample was labeled with TMT-Pro 16 plex (Thermo Scientific) reagents according to the manufacturer’s instructions. The multiplexed sample was cleaned up with a HLB Oasis 1cc (30mg) cartridge. Approximately 60 µg multiplexed peptides were fractionated with high pH reversed-phase C18 UPLC using a 0.5 mm X 200 mm custom-packed UChrom C18 1.8 µm 120Å (nanolcms) column with mobile phases 0.1% (v/v) aqueous ammonia, pH10 in water and acetonitrile (ACN). Peptides were gradient eluted at 20 µL/minute from 2 to 50% ACN in 50 minutes concatenating for a total of 12 fractions using a Waters M-class UPLC (Waters). Peptide fractions were then dried in a speedvac vacuum centrifuge and stored at -20°C until analysis.

##### Mass Spectrometry Analysis

High pH peptide fractions were suspended in 3% (v/v) ACN, 0.1% (v/v) trifluoroacetic acid (TFA) and approximately 1 µg tryptic peptides were directly injected onto a reversed-phase C18 1.7 µm, 130 Å, 75 mm X 250 mm M-class column (Waters), using an Ultimate 3000 nanoUPLC (Thermos Scientific). Peptides were eluted at 300 nL/minute with a gradient from 4% to 25% ACN over 120 minutes then to 40% ACN in 5 minutes and detected using a Q-Exactive HF-X mass spectrometer (Thermo Scientific). Precursor mass spectra (MS1) were acquired at a resolution of 120,000 from 350 to 1500 m/z with an automatic gain control (AGC) target of 3E6 and a maximum injection time of 50 milliseconds. Precursor peptide ion isolation width for MS2 fragment scans was 0.7 m/z with a 0.2 m/z offset, and the top 15 most intense ions were sequenced. All MS2 spectra were acquired at a resolution of 45,000 with higher energy collision dissociation (HCD) at 30% normalized collision energy. An AGC target of 1E5 and 120 milliseconds maximum injection time was used. Dynamic exclusion was set for 20 seconds. Rawfiles were searched against the Human database using MaxQuant v.2.0.3.0. Cysteine carbamidomethylation was considered a fixed modification, while methionine oxidation and protein N-terminal acetylation were searched as variable modifications. All peptide and protein identifications were thresholded at a 1% false discovery rate (FDR). Reporter ion intensities were Cyclic loess normalized and log2 fold changes and p-values were calculated with limma using an R-script.

##### Data analysis

Changes in protein abundance were determined by the “intensity_120mSY5609_._intensity_120mDMSO._lfc” values, describing the log2 fold change in peptide intensity between treated and control samples. Significance was determined by the “intensity_120mSY5609_._intensity_120mDMSO._Pval” column, describing the unadjusted p-value for the change in peptide intensity. Data was plotted in R (3.6.0) with ggplot2 (3.4.4) and pheatmap (1.0.12). Heatmap clustering was automatically generated by the pheatmap function. Rank files for GSEA were generated based on the magnitude, direction, and significance of change. GSEA was run with GSEA v4.3.2 and the gene ontology biological processes geneset (c5.go.bp.v2023.2.Hs.symbols.gmt).

#### Immunofluorescence and western blots

##### Cell Culture

HCT116 (ATCC, CCL-247) cells were cultured in McCoy’s 5A medium (Gibco, 16600082) supplemented with 10% fetal bovine serum (Peak Serum Inc, PS-FB3) and 1% Penicillin-Streptomycin (Gibco, 15140-163) and grown on 150 mm^2^ dishes. Cells were cultured at 37°C with 5% CO_2_.

##### Immunofluorescence

The method used was adapted from Arora et al.^63^ A glass bottom 96 well-plate was treated with collagen (Advanced BioMatrix, 5005-100ML) for 1 hour, then HCT116 cells were seeded on the 96 well glass bottom plate, with 3 wells per condition. Cells recovered overnight. HCT116 cells were treated with either SY-5609 (50 nM) or DMSO. Treatment was stopped after 30 minutes and 6 hours. Cells were fixed with 4% paraformaldehyde for 12 minutes and then washed three times with 1X PBS. Permeabilization was carried out with 0.1% Triton-X in PBS for 20 min at 4°C, then washed three times with 1X PBS. Cells were then incubated in 3% bovine serum albumin (BSA) in PBS for 1 hour at room temperature. Primary antibodies were diluted 1:250 in 3% BSA in PBS and then incubated overnight at 4°C. The following antibodies were used: phospho-CTD ser-5 (Millipore Sigma clone 3E8 04-1572) and RNA Polymerase II Subunit B2 (GeneTex GTX102535). Cells were washed three times with 1X PBS, and secondary antibody conjugated to Alexa Flour 488 goat anti-rabbit (Invitrogen, A78953) or Alexa Fluor 647 goat anti-rat (Invitrogen, A21247) were diluted, 1:1000 in 3% BSA, and incubated for 1 hour at room temperature. DNA content was measured using Hoechst stain, 1:10,000 in PBS, for 5 min at room temperature. Cells were washed twice with 1X PBS, and 1X PBS was added to each well for imaging. Images were captured with a Nikon Eclipse Ti microscope with a 10X 0.5 NA objective. Plots were created using custom MATLAB scripts.

##### Western blots

HCT116 cells were treated with SY-5609 (50 nM) or vehicle (DMSO) and harvested at 30, 75, 180, and 360 minutes. Cells were treated and harvested in biological and technical duplicate. All buffers were supplemented with protease Inhibitors: 1 mM Benzamidine (Sigma-Aldrich, B6506-100G), 1 mM Sodium Metabisulfite (Sigma-Aldrich, 255556-500G), 1 μg/mL Aprotinin (Sigma-Aldrich, A6279-10ML), and 25 mM phenylmethylsulfonyl fluoride (Millipore Sigma, 52332-25GM) immediately before use. Cells were spun at 1,000 rpm, at 4°C for 5 minutes. The supernatant was removed, and the cell pellet was then resuspended in wash buffer (50 mM HEPES pH=7.4, 150 mM NaCl) and spun at 1,000 rpm at 4°C for 5 minutes. The supernatant was removed, and the mass of the cell pellet was measured. The cell pellet was then resuspended in 5:1 (weight:volume) of lysis buffer (1% glycerol, 50 mM HEPES pH=7.4, 150 mM NaCl, 0.5% NP-40, 1 mM EDTA) and nutated at 4°C for 1 hour. The lysate was then spun at 2,300 rpm at 4°C for 10 minutes. The cytoplasmic fraction was removed and snap-frozen in liquid nitrogen and stored at -80°C. RIPA lysis buffer (Thermo Scientific, 89900), supplemented with Benzonase® endonuclease (Millipore Sigma, 101697), was used to resuspend the nuclei. The nuclei were nutated for 1 hour at 4°C. The nuclear extract was then sonicated in a bath sonicator 3 times for 30 seconds, with 1 minute recovery on ice between each sonication. The nuclear extract was then spun at 13,200 rpm at 4°C for 30 minutes. The nuclear extract was aliquoted and then snap-frozen in liquid nitrogen and stored at -80°C. Protein samples were prepared by adding urea (to 2M final concentration), and an equal amount of 2X Loading Buffer. Samples were then boiled at 98°C for 5 minutes and 8 µg of protein per sample were run on an SDS-acrylamide gel, followed by transfer to a nitrocellulose membrane. Post-transfer membranes were stained using Revert 700 Total Protein Stain (Li-Cor, 926-11021) to visualize total protein. Blocking was performed with 5% bovine serum albumin in TBS-T. The following antibodies were used: CTD phospho-serine 5 (Millipore Sigma clone 3E8 04-1572), phospho-CTD ser-7 (Millipore Sigma, clone 4E12 04-1570), phospho-CTD ser-2 (Millipore Sigma clone 3E10 04-1571). HRP-conjugated anti-rat (1:20,000, Invitrogen, 31470) and HRP-conjugated anti-rabbit (1:20,000, Invitrogen, 31460) secondary antibodies were used for detection. Protein was visualized using SuperSignal West Pico PLUS Chemiluminescent Substrate (Fisher Scientific, PI34580) on an Amersham ImageQuant 800. ImageQuant TL 10.2 analysis software was used to quantify and normalize intensity values. Intensity values were normalized to total protein stain.

#### Chromatin Immunoprecipitation Sequencing (ChIP-seq)

##### Cell culture

HCT116 cells (ATCC# CCL-247) were grown to 70-90% confluency in McCoy’s 5A medium and supplemented with 10% Fetal Bovine Serum and 1% Penicillin/Streptomycin antibiotic at 37°C and 5% CO2. Cells were treated with either 10ng/mL IFNγ and DMSO (control) or 10ng/mL IFNγ and 50nM SY-5609. Cells were pre-treated with SY-5609 for 30 minutes followed by IFNγ for 45minutes (or no IFN controls) prior to crosslinking (75 minutes total).

##### Crosslinking and nuclei isolation

Cells were crosslinked with a final concentration of 1% methanol-free formaldehyde for 10 minutes while shaking at 145 RPM on an orbital shaker. Cells were quenched with 2.5M Glycine to a final concentration of 125 mM for 8-10 minutes. Cells were harvested, washed with ice cold PBS (no calcium, no magnesium), and incubated in NRO buffer (10mM Tris-HCl pH8.0, 10mM NaCl, 4mM MgCl_2_, 0.5% NP-40) at a concentration of 80 μL/1 million cells for five minutes twice. Nuclei were pelleted, frozen in liquid nitrogen, and stored at -80°C for future use.

##### IP prep

50 μL of Protein G Dynabeads/reaction was aliquoted into two tubes. Beads were washed with 1 mL Blocking Buffer (1x Phosphate-buffered saline, 0.5% BSA) three times, then resuspended in 250 μL of blocking buffer/reaction. 6 μg/reaction of desired antibody was allowed to bind to the beads for 10 minutes at room temperature with nutation, followed by nutation at 4°C for 3-6 hours. The following antibodies were used: RNAPII Subunit B2 (GeneTex GTX102535), RNAPII CTD phospho-serine 5 (Millipore Sigma clone 3E8 04-1572), and RNAPII CTD phospho-serine 7 (Millipore Sigma, clone 4E12 04-1570).

##### Chromatin shearing

Nuclei were thawed on ice for 1.5-3 hours. Once pellets thawed, nuclei were resuspended in 1 mL Covaris Lysis Buffer B + 1x Protease Inhibitors and incubated for 10 minutes at 4°C with nutation. Nuclei were then washed with 1 mL Covaris Wash Buffer C + 1x Protease Inhibitors and incubated for 10 minutes at 4°C with nutation. Nuclei were washed with 500 μL Shearing Buffer D3 + 1x Protease Inhibitors and resuspended in 1 mL of the same solution per ∼20 million cells. Nuclei were then aliquoted into AFA millitubes (1 mL/tube) and sheared on the Covaris M220 Focused Ultrasonicator for 12 minutes at 6°C, duty factor 10%, peak power 75, 200 cycles/burst. Sheared chromatin was divided into 750-780 μL aliquots (∼15 million cells/aliquot) into pre-chilled Eppendorf tubes, frozen in liquid nitrogen, and stored at -80°C for future use if not immediately proceeding to the IP step.

##### IP conditions

If needed, sheared chromatin was thawed on ice for 2-3 hours. Antibody-bound beads were resuspended in 500 μL cold blocking buffer/reaction, followed by resuspension in the same volume of IP buffer/reaction. Beads were then resuspended in 50 μL IP buffer/reaction, and each reaction (50 μL beads) was aliquoted into a single 2 mL tube. 275 μL IP buffer (15mM Tris-HCl pH8.0, 150mM NaCl, 1mM EDTA, 1% Triton X-100) was added to each tube, followed by 500 μL IP concentrate, and ∼360 μL of sample. This means that each aliquot of approximately 720 μL of sheared chromatin from roughly 15 million cells was split into two tubes at the IP step. This volume is slightly less than that which was aliquoted during the shearing step to account for input volume and loss of volume during pipetting steps. Splitting each IP into two tubes was necessary to dilute the amount of SDS present in the IP mixture to a final concentration of 0.03% to maximize antibody-protein binding. IPs were nutated at 4°C for 12-18 hours. 10 μL of sheared chromatin from each treatment condition per IP was aliquoted into a separate tube and stored at -20°C to provide an input sample.

##### Washing and reversing crosslinks

Beads were washed with IP Buffer, RIPA wash buffer (20mM Tris HCl pH8.0, 500mM NaCl, 1mM EDTA, 1% Triton X-100, 0.1% SDS), LiCl Wash Buffer (20mM Tris HCl pH8.0, 500mM NaCl, 1mM EDTA, 1% sodium deoxycholate, 1% NP-40), and TE Salt Buffer (10mM Tris HCl pH8.0, 1mM EDTA, 50mM NaCl) two times each to remove non-specific protein binding interactions. All washes were performed on ice and incubated for 50-80 seconds. Following the final wash, beads were resuspended in 200 μL of Proteinase K Buffer containing 5 μL of 20 mg/mL Proteinase K. Beads were incubated on a Boekel Groovin’ Tubes Thermomixer set to 50°C mixing at 750 RPM for one hour, or on a 50°C heat block and manually resuspended every 20 minutes. Beads were then incubated at 65°C for one hour. 190-195 μL of supernatant was removed from the beads and transferred to 0.2 mL tubes. Input chromatin was thawed and brought to 200 μL by adding 190 μL of aforementioned Proteinase K mix. All inputs and IPs were incubated at 65°C for 17.5 hours.

##### DNA isolation and precipitation

Samples were vortexed with an equal volume of phenol:chloroform:isoamyl (25:24:1) to generate aqueous and organic phases. Both phases were added to a phase lock tube and nutated for 10 minutes at room temperature. Phases were separated via centrifugation (Phase Lock tubes contain a gel that forms a barrier between phases) and an equal volume of either phenol:chloroform:isoamyl or pure chloroform was added to the aqueous phase to perform a second DNA extraction. Phase lock tubes were again nutated for 10 minutes at room temperature and centrifuged to isolate the aqueous phase. The aqueous phase was moved to a fresh tube, to which 20 μg GlycoBlue, 1/10 sample volume of 3M NaOAc pH 5.2, 10 mM MgCl_2_, and 2.5x total volume of 100% ethanol was added. Tubes were mixed via vortexing and incubated at -20°C for one to two days. DNA pellets were isolated via 30 minute centrifugation at 12,000 x g. Supernatant was removed, and pellets were washed with 750 μL of 75% ethanol. Ethanol was removed and pellets were air-dried for 5-12 minutes. Once dry, pellets were suspended in 20-30 μL of 0.1x TE Buffer. DNA concentrations were determined via Qubit and samples were stored at -20°C.

##### Library preparation

Libraries were prepared using the KAPA Hyperprep Kit. 1 ng of total DNA from each sample was diluted in 50 μL 0.1x TE buffer and subjected to end repair and A-tailing. Adapter ligation was performed using an adapter stock concentration of 300 nM per manufacturer’s instructions and was allowed to proceed for one hour at 20°C. Following ligation, a 0.8x bead cleanup was performed, and the libraries were amplified according to manufacturer’s instructions. Samples were amplified for 14 cycles. Following amplification, libraries were subjected to two consecutive 1x bead-based cleanups to remove protein contaminants and adapter dimers. Libraries were quantified via Qubit and imaged using capillary electrophoresis on an Agilent 2200 Tapestation to identify average size. Libraries that exhibited overamplification were subjected to a reconditioning PCR, in which 16-19 μL of library was mixed with a proportional volume of primers and PCR mastermix and subjected to one cycle of amplification and a subsequent 1x bead based cleanup. This served to resolve daisy-chained DNA fragments, and reconditioned libraries exhibited the anticipated size peaks on the tapestation.

Libraries were pooled by mass and concentration according to a 5:3:1 ratio of total RNAPII (NTD): phosphomarks:input samples. Library pools were subsequently diluted to 10 nM in 30 μL according to sequencing core instructions. SY-5609 biological replicates were sequenced to a total requested depth of 600 million paired-end reads. All samples were sequenced on a NovaSeq 6000 by the Gates Institute Genomics Core at CU Anschutz.

##### Data processing

Initial sample processing and quality control was performed using the Dowell Lab ChIP-Flow pipeline (https://github.com/Dowell-Lab/ChIP-Flow, v19.10.0). If needed, corrupted raw data files were amended using the bbmap repair function (v38.05). QC metrics showed one biological replicate was higher quality in each condition and therefore only one replicate was used for subsequent analyses. We justify including the results because we do not make novel claims from the data, and the results are supportive of other experiments and consistent with other studies that used CDK7 inhibitors. Briefly, read quality was assessed using FastQC (v0.11.8), and adapters were trimmed from reads via bbmap. Reads were mapped to the GRCh38 reference genome (UCSC Assembly, December 2013) using hisat2 (v2.1.0). Library complexity was estimated using preseq (v2.0.3) and compressed into .tdf files for genome browser visualization using IGVTools (v2.3.75). FastQC, preseq, and rseqc (v 3.0.0) metrics were compiled into a single ouput via MultiQC (v1.15). Duplicate, multimapping, and non-mapping reads were removed via sambamba. Indices for bams containing only uniquely mapped reads were generated using samtools 1.18.0. Data was then normalized to 1x read coverage via the DeepTools (v3.5.2) bamCoverage package. hg38 blacklist regions were excluded from analysis. The resulting bigwig files were used to create a matrix using DeepTools computeMatrix to calculate scores for each sample at desired genomic regions. These matrices were plotted using DeepTools plotProfile package; bin size = 50 bp.

##### Metagene Analysis

5’ and 3’ metagenes from HCT116 ChIP-seq (GEO accession GSE268531) were generated almost identically to PRO-seq using the filtered highly expressed gene set (n=453), but representing one replicate rather than two. Counting differed between the two protocols, in that midpoints of valid fragments were counted in a non-strand-specific manner for ChIP-seq data rather than the 5’-most base as with PRO-seq data.

##### Pause index analysis

Pause indexes were calculated by splitting Refseq genes into two annotation files: one delineating 401bp regions around the annotated TSS (–100 to +300 relative to TSS) and the other specifying the gene body (+301 to polyA site). Reads were then counted within those two regions, discarding any along boundaries, using the featurecounts function of Rsubread (version 2.12.3, R version 3.6.1) with the following parameters: useMetaFeatures=FALSE, allowMultiOverlap=TRUE, countMultiMappingReads=FALSE, fracOverlap=1, strandSpecific=1. The counts were then run through a custom python script (python version 3.6.3) wherein 3975 genes with the highest count sums across samples were isolated and a length-normalized read count ratio was calculated between the pause region and the rest of the gene.

##### Plotting

Plotting was done in R (version 4.2.3) with the packages ggplot2 (version 3.4.3), dplyr (version 1.1.3), reshape2 (version 1.4.4) and cowplot (version 1.1.1).

#### 3’-end Metagene analyses for TT-seq and mNET-seq (Velychko et al. 2024)

Published data^16^ were downloaded from the Short Read Archive (Parent GEO accession GSE218269, TT-seq SRA sample accessions SRR22327405-SRR22327416; NET-seq SRA sample accessions SRR22327385-SRR22327391). Reads were trimmed using bbduk (bbmap v38.05) and mapped with hisat2 (v2.1.0) and samtools (v1.10). 3’ metagenes were generated as in PRO-seq with the following differences: The gene set used was derived from all Refseq gene annotations (n=41388), and as the samples were paired end, the 5’ most base of the first-in-pair read was counted.

##### Plotting

Plotting was done in R (version 4.2.3) with the packages ggplot2 (version 3.4.3), dplyr (version 1.1.3), reshape2 (version 1.4.4) and cowplot (version 1.1.1).

#### Phosphoproteomics sample preparation and phospho-enrichment

HCT116 and OV90 cells treated for 1 hour with the CDK7 inhibitor SY-5609 or vehicle (DMSO) were immediately harvested with 5% (w/v) sodium dodecyl sulfate (SDS), 10 mM tris(2-carboxyethylphosphine) (TCEP), 40 mM 2-chloroacetamide and 50 mM Tris-HCl, pH 8.5 with protease and phosphatase inhibitors (Roche) then boiled for 10 minutes. Cell lysates were probe-sonicated and stored at -75°C. Approximately 600 to 900 µg protein lysates were LysC/trypsin digested using the SP3 method.^59^ Briefly, 1000 µg carboxylate-functionalized speedbeads (Cytiva Life Sciences) were added followed by the addition of acetonitrile (ACN) to 80% (v/v) inducing binding to the beads. The beads were washed twice with 80% (v/v) ethanol and twice with 100% acetonitrile. Proteins were digested in 50 mM Tris-HCl, pH 8.5, with Lys-C/Trypsin (Promega) incubating at 37°C shaking at 2000rpm overnight. Tryptic peptides were desalted using Waters 6cc (150mg) HLB Oasis cartridges according to the manufacturer’s instructions and dried in a speedvac vacuum centrifuge and stored at -20°C. Tryptic peptides were phosphoenriched using the High-Select TiO_2_ phosphoenrichment kit (Thermo Scientific). Samples were suspended in the Binding/Equilibration buffer, loaded onto the TiO_2_ column, washed and eluted. The elution and the unretained flow through fractions were dried immediately in a speedvac vacuum centrifuge. The TiO_2_ elution was immediately resuspended with 1% TFA, 3% ACN and stored at 4°C, while the unretained flowthrough fraction was subjected to a sequential phosphoenrichment with the High-Select Fe-NTA phosphoenrichment kit (Thermo Scientific). The TiO_2_ unretained flow through fraction was suspended in the Fe-NTA Binding/Wash Buffer, loaded onto the Fe-NTA column, washed and eluted. The Fe-NTA elution was immediately resuspended with 1% TFA, 3% ACN and stored at 4°C. The TiO_2_ and the Fe-NTA elutions were combined and cleaned up using a Waters M-class UPLC with a PDA detector and a custom fabricated 0.5 mm X 180 mm rpC18 1.9 µm 120 Å column (Dr Maisch Gmbh) column with a gradient from 2% to 50% ACN in 10 minutes then dried immediately in a speedvac vacuum centrifuge. Peptides were then labeled with TMT 10 plex reagents (Thermo Scientific) according to the manufacturer’s instructions. The multiplexed sample was immediately cleaned up with a HLB Oasis 1cc (30mg) cartridge and dried in a speedvac vacuum centrifuge. Multiplexed phosphopeptides were then fractionated on a rpC18 column with high pH mobile phases (0.1% ammonium hydroxide) using a Waters M-class UPLC with a PDA detector and a custom-fabricated 0.5 mm X 180 mm rpC18 1.9 µm 120 Å column (Dr Maisch Gmbh) column with a gradient from 2% to 20% ACN in 50 minutes. Thirty second fractions were concatenated for a total of 12 phosphoproteome fractions. High pH fractions were immediately dried in a speedvac vacuum centrifuge and stored at -20°C until LC/MS analysis.

#### Phosphoproteomics mass spectrometry and data analyses

High pH fractionated phosphopeptides were suspended in 3% (v/v) acetonitrile (ACN), 0.1% (v/v) trifluoroacetic acid (TFA) and directly injected onto a reversed-phase CSH rpC18 1.7 µm, 130 Å, 75 mm X 250 mm M-class column (Waters), using a Vanquish Neo nanoUPLC (Thermo Scientific). Phosphopeptides were eluted at 300 nL/minute with a gradient from 4% to 16% ACN, 0.1% (v/v) formic acid in 120 minutes then to 25% ACN in 5 minutes and detected using a Orbitrap Exploris 480 mass spectrometer (Thermo Scientific). Precursor mass spectra (MS1) were acquired at a resolution of 120,000 from 350 to 1500 m/z with a standard automatic gain control (AGC) target and Auto maximum injection time mode. Precursor peptide ion isolation width for MS2 fragment scans was 0.7 m/z with a 3 second cycle time. All MS2 spectra were acquired at a resolution of 15,000 using TMT Turbo with higher energy collision dissociation (HCD) at 34% normalized collision energy. A custom normalized AGC target of 200% and Auto maximum injection time was used. Dynamic exclusion was set for 30 seconds. Rawfiles were searched against the Uniprot Human database UP000005640 downloaded 08/26/2022 using MaxQuant v2.6.3.0. Cysteine carbamidomethylation was considered a fixed modification, while methionine oxidation, protein N-terminal acetylation and phosphorylation at serine, threonine or tyrosine were searched as variable modifications. All identifications were thresholded at a 1% false discovery rate (FDR). TMT reporter ion intensities were normalized using the cyclic loess method, then differential phospho-occupancies were calculated using Limma (Bioconductor.org) to generate a linear model for fold-change estimation and standard error with an Empirical Bayes implementation of an independent t-test where the null distribution is inferred from the global data, and treatment distribution is Bayesian updated. Adjusted p-values were then generated using the Benjamini-Hochberg limma interpretation.

