## Supplemental Information for "Multi-omics and biochemical reconstitution reveal CDK7-dependent mechanisms controlling RNA polymerase II function at gene 5’- and 3’-ends"

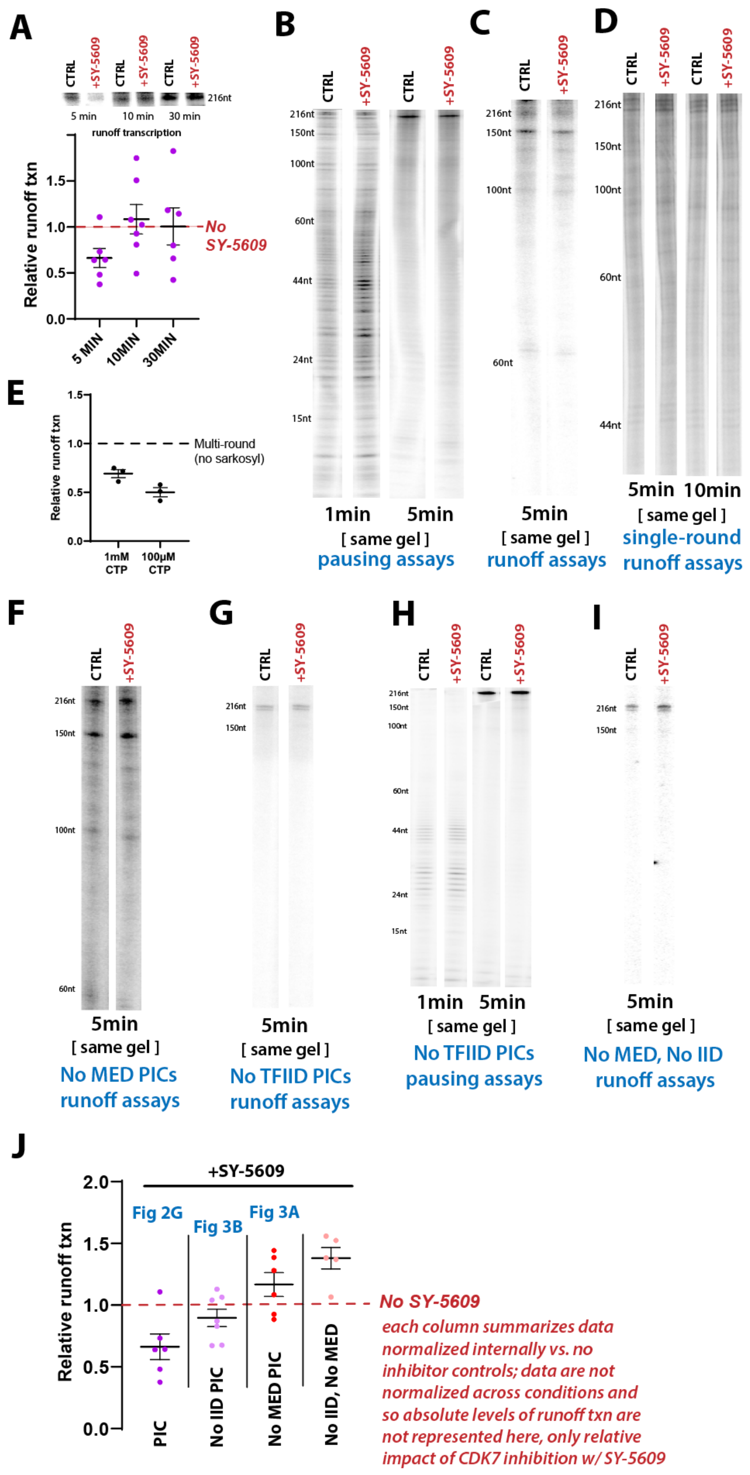


**Figure S1. Representative gel images** (related to Figure 2 & Figure 3)
(A) Plot showing quantitation of RNAPII runoff transcription +SY-5609, normalized to no inhibitor controls. The CDK7-dependent effects are lost at longer timepoints. Because we were interested in identifying CDK7-dependent mechanisms associated with RNAPII initiation, the relevant timeframe was within 5 minutes NTP addition. We hypothesize that PIC stabilization occurs in CDK7-inhibited conditions, based upon evidence that PIC disassembly is sensitive to CDK7 kinase activity in yeast and human cells.[1-5] This potential "secondary effect" may contribute to equalizing runoff transcription levels over time, as well as low levels of CDK7 activity that occur over longer timeframes (e.g. in vitro kinase assays, **Figure 1F, G**). Top: representative runoff transcription data after 5, 10, or 30 minutes ±SY-5609.

(B-D; F-H) Representative gel images from *in vitro* transcription assays shown in **Figure 2** and **Figure 3**. Each timepoint was imaged at the same exposure and contrast ±SY-5609, across the entire lane; this uniform exposure and contrast was used for data quantitation shown throughout the paper. Gel composition was 7%, 12%, or 18% acrylamide depending on desired resolution.
(E) Verification that more transcription occurs with multi-round vs. single-round conditions. Quantitation of runoff transcripts from two different experimental conditions (n=3 each; 10min reactions), in which the cold CTP chase was 1mM or 0.1mM. For single-round transcription, sarkosyl was added to a final concentration of 0.2% 1min after NTPs.
(I) Representative lanes from runoff assays with PICs lacking Mediator and TFIID (TBP instead of TFIID), ±SY-5609. These results are included in scatterplot shown in panel H.
(J) Mediator and TFIID effects are additive in ±SY-5609 comparisons. Each column was normalized independently vs. no inhibitor controls (dashed line), but data are shown on a single plot to emphasize trends as a function of Mediator and TFIID +SY-5609. Each column shows data from main text figures (noted in blue font), except for the "No IID, No MED" experiments, which are shown in panel G (n=6, 7, 6, 5 replicates across columns).

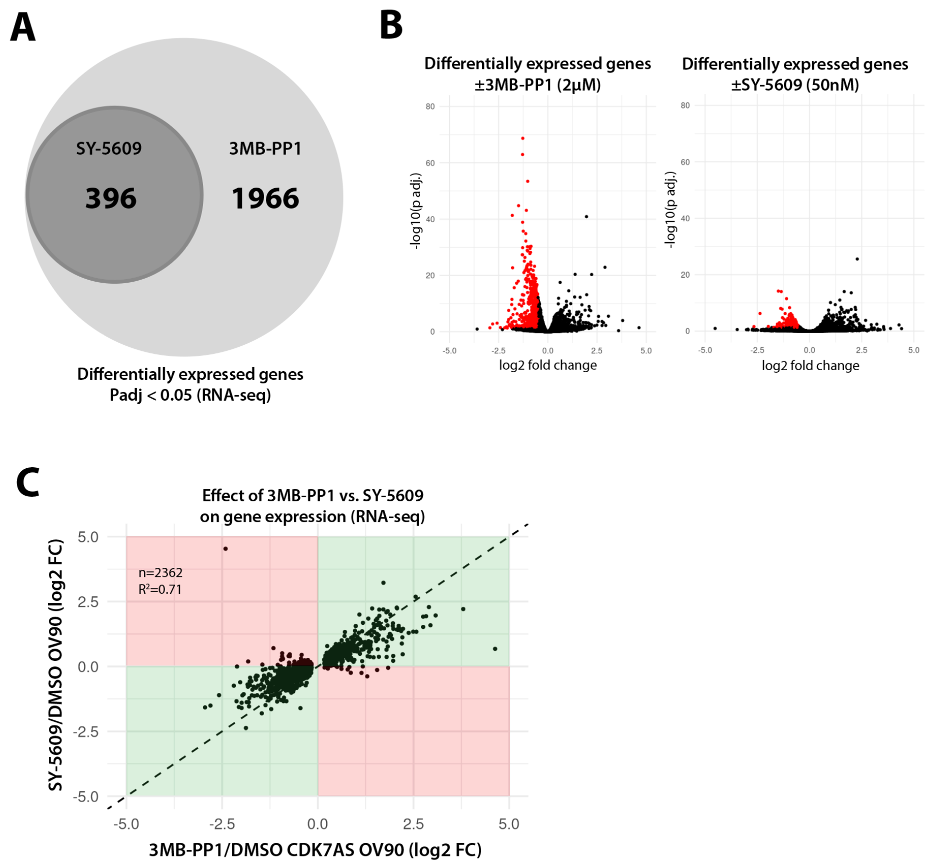


**Figure S2. SY-5609 causes gene expression changes similar to CDK7AS cell line treated with 3MB-PP1 ATP analog** (related to Figure 4)

(A) Venn diagram showing significantly changing genes ±CDK7 inhibitor (Padj < 0.05) in each cell line (OV90 or OV90 CDK7AS).

(B) Volcano plots showing differentially expressed genes (RNA-seq) in CDK7AS OV90 cells ±3MB-PP1 or OV90 cells ±SY-5609. Genes significantly downregulated (Padj < 0.05) are highlighted in red (n=1735 for 3MB-PP1; n=146 for SY-5609).

(C) Scatterplot showing the log2 fold changes in gene expression in CDK7AS OV90 cells treated with 3MB-PP1 (x-axis) compared to SY-5609 (y-axis), each relative to DMSO controls. Each point represents an individual gene; genes significantly changing (Padj < 0.05) with 3MB-PP1 treatment are shown. Genes in the red-shaded quadrants are upregulated in one condition but downregulated in the other, while genes in the green-shaded quadrants show consistent changes (upregulated or downregulated) with either inhibitor (3MB-PP1 or SY-5609), despite the different cell lines and inhibitor concentrations used. The dashed line represents a theoretical perfect fold-change correlation of 1.0 between the two cell lines/treatments. A linear regression line (not shown) with $R^2 = 0.71$ indicates a high degree of correlation in gene expression changes between the two conditions (n=2362 genes).

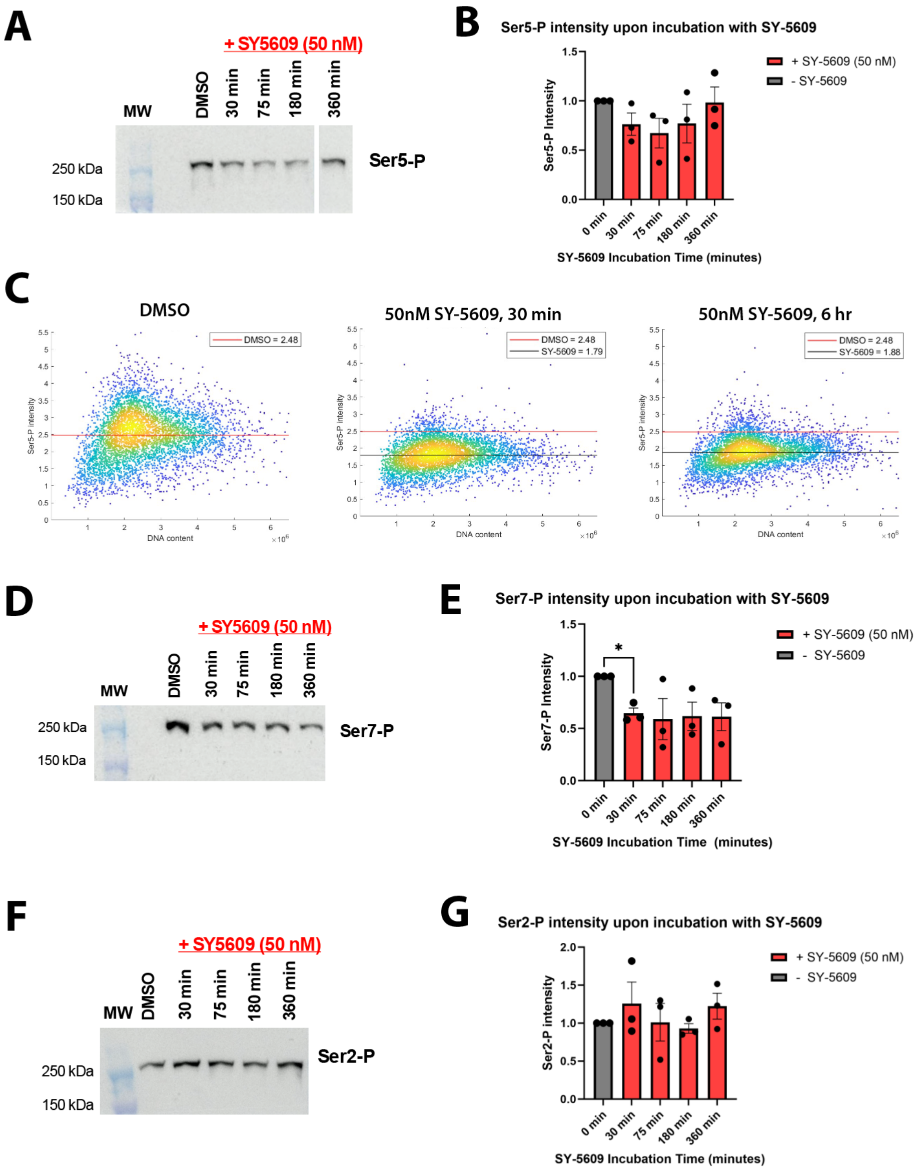


**Figure S3. SY-5609 causes global decreases in RNAPII CTD phosphorylation levels** (related to Figure 4)

(A) Representative western blot data showing reduced ser5-P levels from nuclear extracts of SY-5609-treated cells.

(B) Quantitation of western blot data from 3 experiments (2 biological replicate samples +1 technical replicate). Total protein and a total protein stain were used as a loading control.

(C) Single-cell immunofluorescence data from untreated (DMSO) or SY-5609-treated cells. Shown are density scatterplots of the ser5-P intensity normalized to the total RNAPII intensity in each cell; each spot represents a single cell. Red line indicates the median of the cell population treated with DMSO, black lines indicate median of cell population treated with SY-5609.

(D) Representative western blot data showing reduced ser7-P levels from nuclear extracts of SY-5609-treated cells.

(E) Quantitation of western blot data from 3 experiments (2 biological replicate samples +1 technical replicate). Total protein and a total protein stain were used as a loading control.

(F) Representative western blot data showing ser2-P levels from nuclear extracts of SY-5609-treated cells.

(G) Quantitation of western blot data from 3 experiments (2 biological replicate samples +1 technical replicate). Total protein and a total protein stain were used as a loading control.

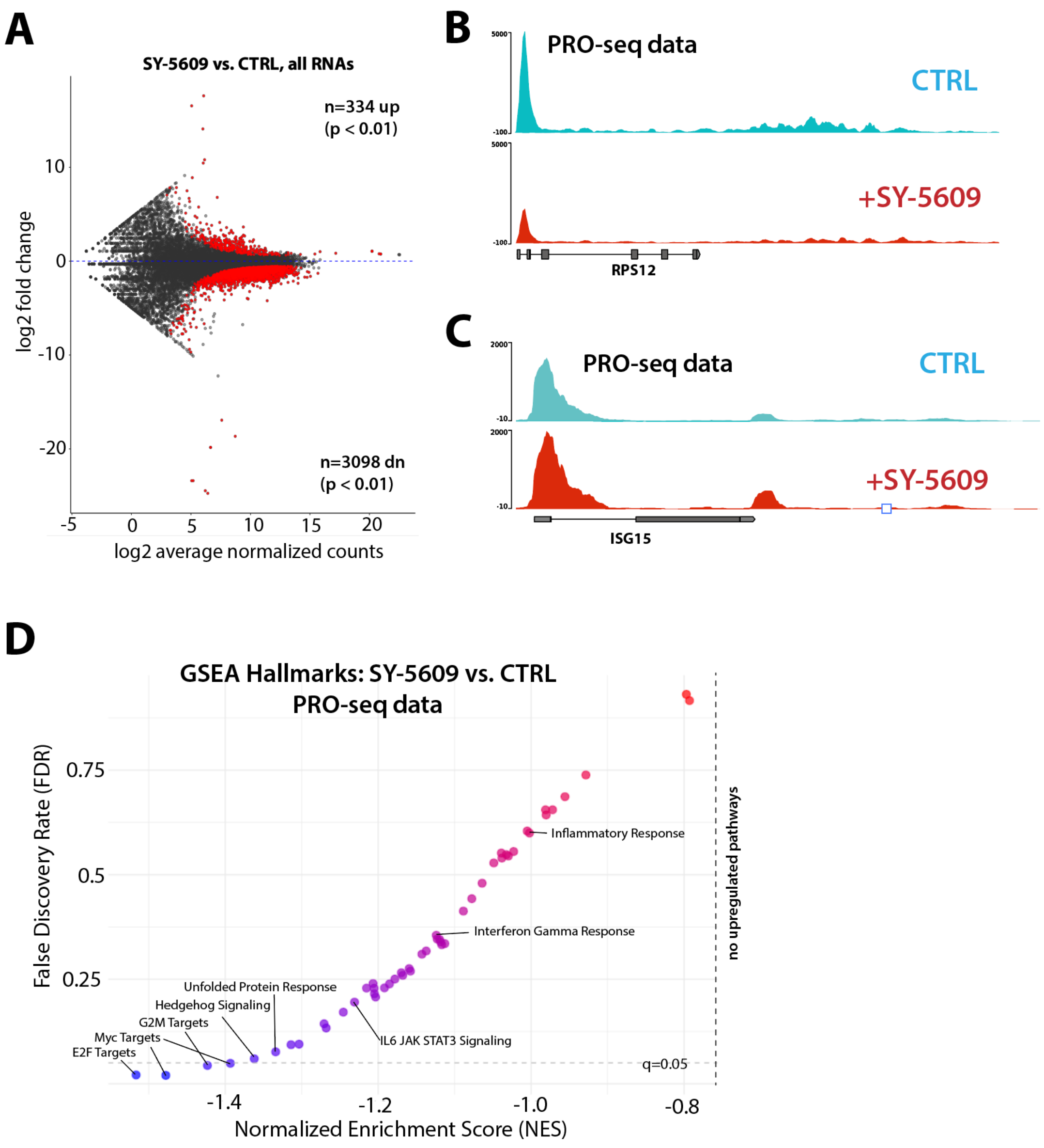


**Figure S4. Overview of gene expression changes in CDK7-inhibited HCT116 cells** (related to Figure 4)

(A) MA plot showing differentially expressed transcripts in SY-5609-treated cells (PRO-seq data). Red dots indicate significantly changing transcripts (p-val < 0.01).

(B) Genome browser traces of PRO-seq reads at a constitutively active gene (RPS12) and

(C) an IFNγ inducible gene (ISG15) in control vs. SY-5609-treated cells.

(D) Summary plot of GSEA results (FDR vs. NES; Hallmarks gene set) showing down-regulated pathways in CDK7-inhibited cells. Note that no pathways showed a positive NES and so the right side of the plot is not shown.

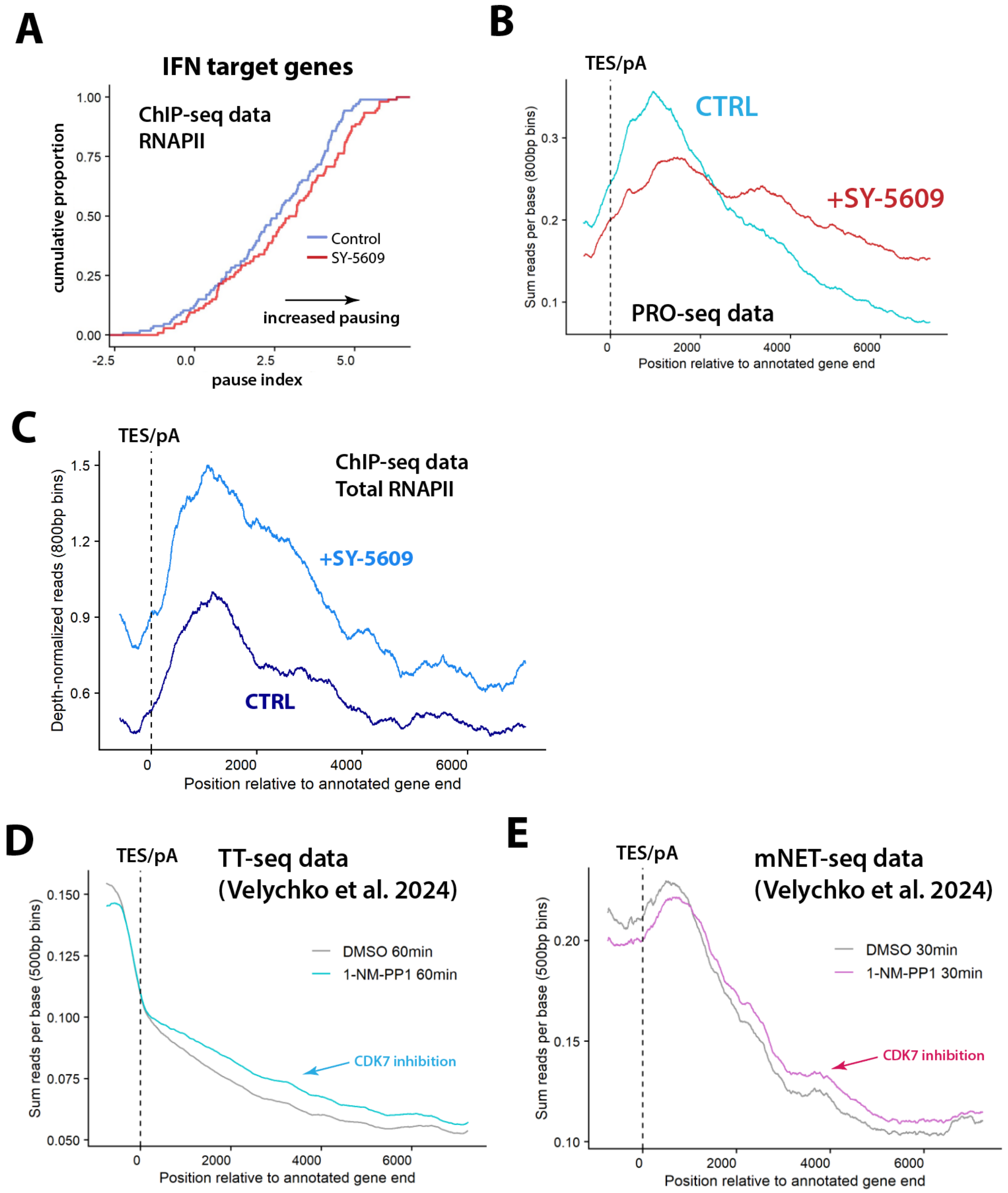


**Figure S5. CDK7 regulates RNAPII function at gene 5'-ends and 3'-ends** (related to Figure 4 & 5)
(A) Pause index cumulative distribution plot from RNAPII ChIP-seq from control and SY-5609 treated cells. Analysis was restricted to IFNγ-stimulated genes (n=96).
(B) Metagene analysis at 3'-ends of highly expressed genes in control and SY-5609 treated conditions (PRO-seq data, n=220). Position 0 indicates the polyA site.
(C) Consistent with PRO-seq results, metagene plot of RNAPII ChIP-seq data shows elevated levels of RNAPII after the polyA site in SY-5609-treated cells compared with untreated controls (n=117). Position 0 indicates the polyA site.
(D) The same metagene analysis used for panel B was applied to TT-seq data from Velychko et al.[3] in CDK7AS cells. CDK7 inhibition (NM-PP1 analog) increased 3'-readthrough transcription (n=11570).
(E) Metagene analysis also applied to mNET-seq data from Velychko et al.[3] in CDK7AS cells (n=11299), again showing 3'-readthrough transcription with CDK7 inhibition, in agreement with our results with SY-5609.

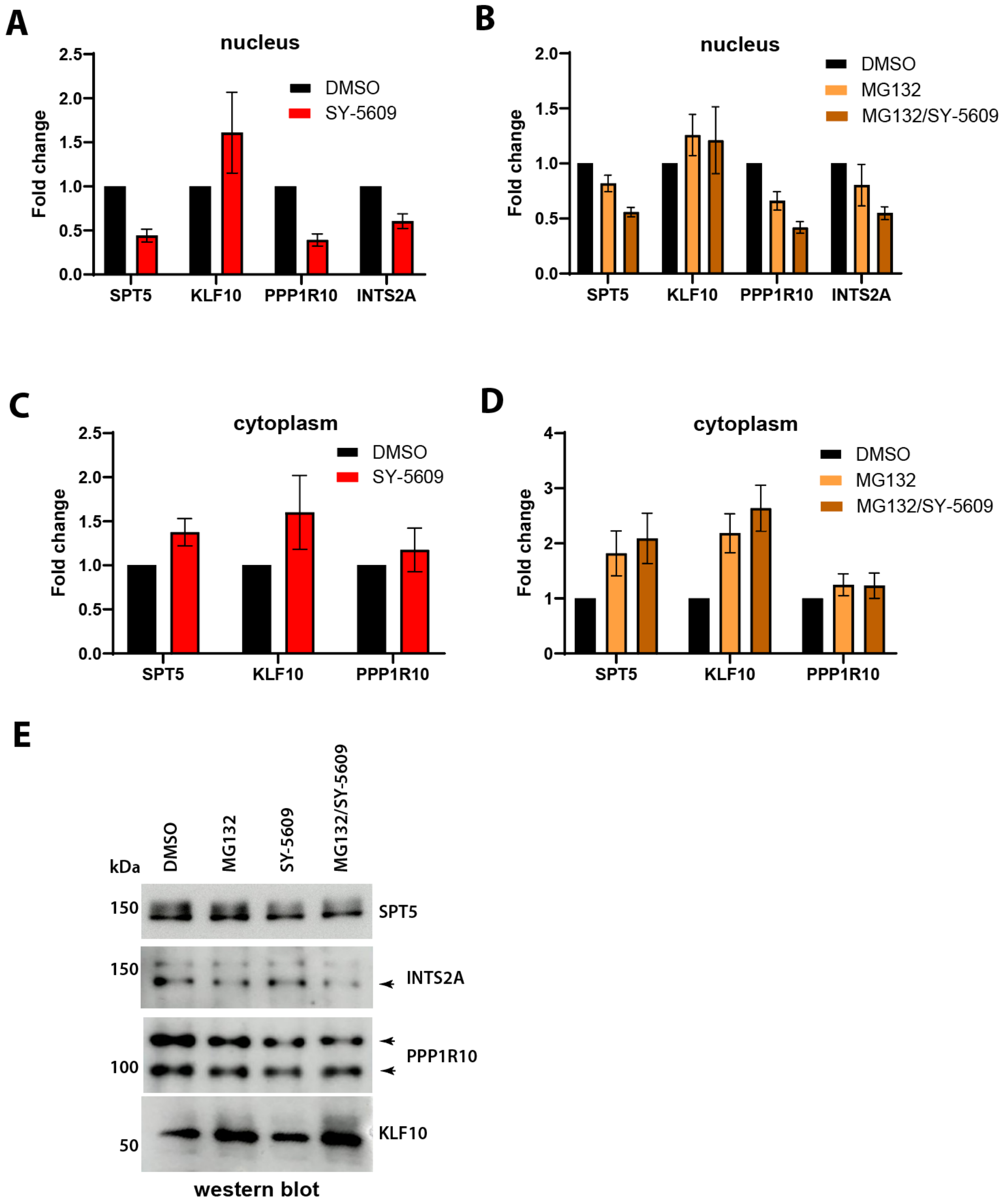


**Figure S6. Quantitative western blots support quantitative MS results** (related to Figure 5 & 6)
(A) Nuclear extracts from HCT116 cells treated with 50nM SY-5609 (or DMSO CTRL; t=2h) were probed for each protein shown; each of these proteins showed decreased levels based on quantitative MS. Total protein and a total protein stain was used as a loading control (n=3-4 for each factor). Bars represent standard error of the mean.
(B) Results from nuclear extracts treated with proteasome inhibitor MG132 (5$\mu$M) or SY-5609/MG132 combination. MG132 had little impact on SPT5, PPP1R10/PNUTS, and INTS2A, suggesting these factors are not degraded.
(C) Cytoplasmic extracts were probed as described in A (n=3-5 for each factor); bars represent standard error of the mean. Levels of SPT5, KLF10, and PNUTS increase in the cytoplasm +SY-5609, suggesting they are exported from the nucleus. Note that INTS2A antibody did not produce reliable data from cytoplasmic extracts.
(D) Cytoplasmic extracts were probed as described in B.
(E) Representative blots from nuclear extract experiments. For INTS2A and PPP1R10, arrows denote bands used for quantitation, which were confirmed based upon RNAi knockdowns.

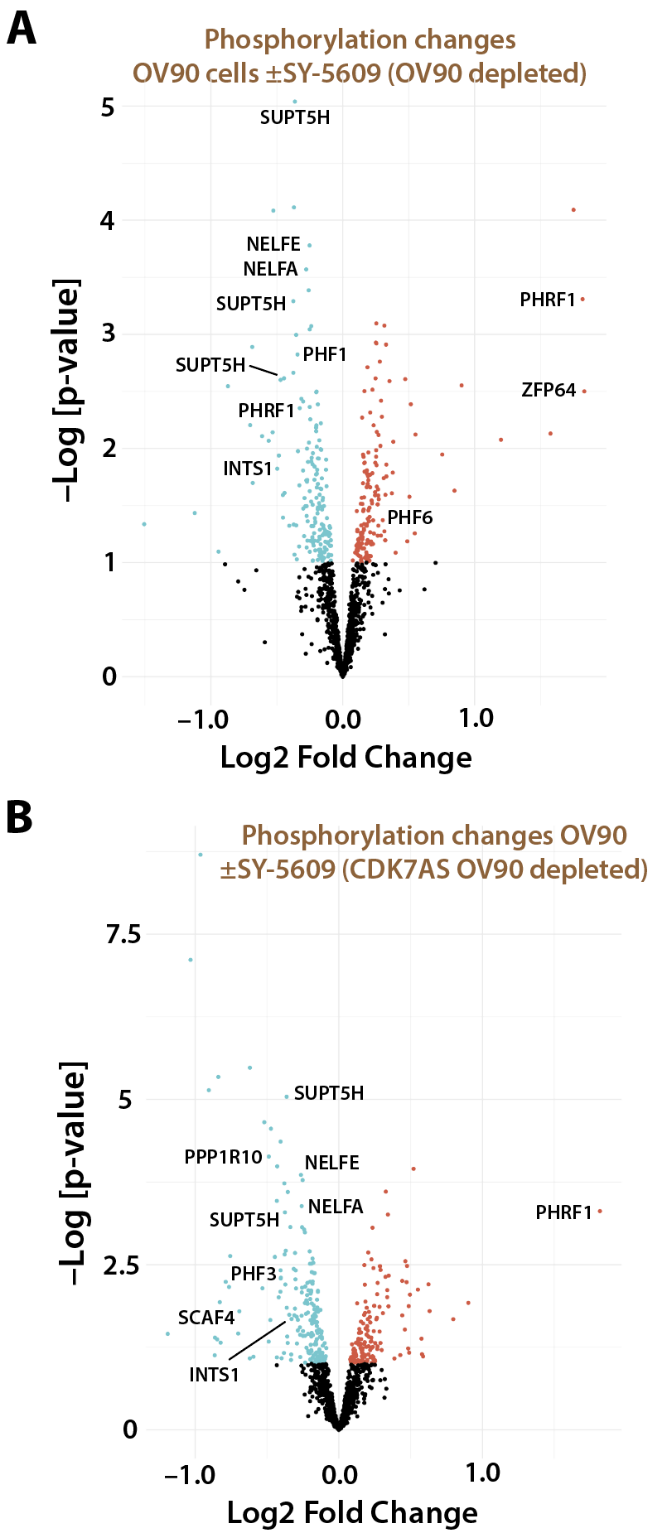


**Figure S7. CDK7 phosphorylates many RNAPII elongation and termination/3'-processing factors (OV90 cells)** (related to Figure 6)

(A) Volcano plot showing proteins with phosphorylation changes ±SY-5609 (50nM, 1h) in OV90 cells. For simplicity, only proteins that were nuclear depleted in OV90 cells +SY-5609 are included here (n=1227). Select proteins are labeled on plot; full list in **Table S6**.

(B) Volcano plot showing proteins with phosphorylation changes ±SY-5609 (50nM, 1h) in OV90 cells. For simplicity, only proteins that were nuclear depleted in OV90 CDK7AS cells +SY-5609 are included here (n=1157). Select proteins are labeled on plot; full list in **Table S6**.

**Supplemental Tables:**

**Table S1.** MS data from immobilized template + nuclear extract experiments.

**Table S2.** Quantitative MS data for HCT116 ±SY-5609; t=30min & t=120min.

**Table S3.** Quantitative MS data for OV90 ±SY-5609; t=30min & t=120min.

**Table S4.** Quantitative MS data for CDK7AS OV90 ±3MB-PP1; t=4h.

**Table S5.** Quantitative phospho-proteomics MS data for HCT116 ±SY-5609; t=60min.

**Table S6.** Quantitative phospho-proteomics MS data for OV90 ±SY-5609; t=60min.
